# Deep sequencing of proteotoxicity modifier genes uncovers a Presenilin-2/beta-amyloid-actin genetic risk module shared among alpha-synucleinopathies

**DOI:** 10.1101/2024.03.03.583145

**Authors:** Sumaiya Nazeen, Xinyuan Wang, Dina Zielinski, Isabel Lam, Erinc Hallacli, Ping Xu, Elizabeth Ethier, Ronya Strom, Camila A. Zanella, Vanitha Nithianandam, Dylan Ritter, Alexander Henderson, Nathalie Saurat, Jalwa Afroz, Andrew Nutter-Upham, Hadar Benyamini, Joseph Copty, Shyamsundar Ravishankar, Autumn Morrow, Jonathan Mitchel, Drew Neavin, Renuka Gupta, Nona Farbehi, Jennifer Grundman, Richard H. Myers, Clemens R. Scherzer, John Q. Trojanowski, Vivianna M. Van Deerlin, Antony A. Cooper, Edward B. Lee, Yaniv Erlich, Susan Lindquist, Jian Peng, Daniel H Geschwind, Joseph Powell, Lorenz Studer, Mel B. Feany, Shamil R. Sunyaev, Vikram Khurana

## Abstract

Whether neurodegenerative diseases linked to misfolding of the same protein share genetic risk drivers or whether different protein-aggregation pathologies in neurodegeneration are mechanistically related remains uncertain. Conventional genetic analyses are underpowered to address these questions. Through careful selection of patients based on protein aggregation phenotype (rather than clinical diagnosis) we can increase statistical power to detect associated variants in a targeted set of genes that modify proteotoxicities. Genetic modifiers of alpha-synuclein (ɑS) and beta-amyloid (Aβ) cytotoxicity in yeast are enriched in risk factors for Parkinson’s disease (PD) and Alzheimer’s disease (AD), respectively. Here, along with known AD/PD risk genes, we deeply sequenced exomes of 430 ɑS/Aβ modifier genes in patients across alpha-synucleinopathies (PD, Lewy body dementia and multiple system atrophy). Beyond known PD genes *GBA1* and *LRRK2*, rare variants AD genes (*CD33*, *CR1* and *PSEN2*) and Aβ toxicity modifiers involved in RhoA/actin cytoskeleton regulation (*ARGHEF1, ARHGEF28, MICAL3, PASK, PKN2, PSEN2*) were shared risk factors across synucleinopathies. Actin pathology occurred in iPSC synucleinopathy models and RhoA downregulation exacerbated ɑS pathology. Even in sporadic PD, the expression of these genes was altered across CNS cell types. Genome-wide CRISPR screens revealed the essentiality of *PSEN2* in both human cortical and dopaminergic neurons, and *PSEN2* mutation carriers exhibited diffuse brainstem and cortical synucleinopathy independent of AD pathology. *PSEN2* contributes to a common-risk signal in PD GWAS and regulates ɑS expression in neurons. Our results identify convergent mechanisms across synucleinopathies, some shared with AD.

## INTRODUCTION

Proteinopathies are age-related diseases in which specific proteins aggregate and form inclusions in distinct cell types. Different neurodegenerative diseases can be associated with aggregation of the same protein. For example, ɑ-synuclein (ɑS)-rich inclusions, including Lewy bodies (LBs) and glial cytoplasmic inclusions (GCIs), are the hallmark pathology of synucleinopathies, including Parkinson’s disease (PD), dementia with Lewy bodies (DLB) and multiple system atrophy (MSA)^1^. Equally, patients not infrequently have mixed pathologies^2,3^. For example, more than 50% of Alzheimer’s disease cases have the classic Lewy bodies^4,5^, and the hallmark β-amyloid (Aβ) and tau pathologies of AD are common in DLB and Parkinson’s disease dementia (PDD), but also found in a significant proportion of PD cases (>30%)^3,6–8^. These findings raise two fundamentally important questions we sought to shed light on in the current investigation: first, to what extent do synucleinopathies share a common underlying biological basis, and second, are there share risk factors among different proteinopathies?

Rare variants in the ɑ-synuclein-encoding gene *SNCA* suggest a shared basis across synucleinopathies. For example, in familial synucleinopathies, individual rare variants lead to a spectrum of possible disease outcomes, from PD to PDD to DLB, within the same family^9^. Two Mendelian synucleinopathies – caused by either the *SNCA* G51>D point mutation or by triplication at the *SNCA* locus– are associated with diffuse ɑS inclusions characteristic of PD and MSA^10,11^. Additional genetic evidence is emerging to support this. For example, patients harboring mutations in the glucocerebrosidase-encoding gene *GBA1* are at elevated risk of clinical PD, PDD, or DLB^12,13^, and recent publications suggest an association of *GBA1* variants also with MSA. At a cellular level, distinct synucleinopathies could result from the differential vulnerability of distinct brain regions and circuits to ɑS pathology^14^. At a biophysical level, they have been associated with distinct pathologic amyloid conformers, or “strains”^15–19^. It is thus plausible that different clinical presentations, even in patients with identical primary mutations (for example, in *GBA1* or *SNCA*), could stem from other environmental factors or genetic background modifiers that lead to either distinct ɑS conformational states or glio-neuronal vulnerability patterns (or both).

The presence of variable pathologies in most patients with neurodegenerative diseases may reflect the coexistence of different disease processes in the same patient or potentially a shared mechanistic link among different proteinopathies. Moreover, a potential shared link may depend on the existence of overt proteinaceous aggregates. Genetic, transcriptomic, and proteomic interrogations of tractable cellular and organismal models to answer this question have yielded conflicting results^20^. However, evidence for the overlapping genetic basis of neurodegenerative and neuropsychiatric diseases, including synucleinopathies is emerging from multiple recent human genetic studies^21–23^. For example, coding variants at the tau-encoding *MAPT* locus lead to frontotemporal dementia and parkinsonism (a tauopathy)^24^. Still, single nucleotide polymorphisms (SNPs) in linkage with *MAPT* (and specifically the tau H1 haplotype) are among the best-validated risk factors for PD with growing evidence in AD also^25^. The PD association holds in pathologically confirmed synucleinopathy cases and does not seemingly hinge on overt tau aggregation ^26^. In other examples, the *APOE, TMEM175* and *BIN1* risk factors suggest crossover risk between AD and DLB^13,21,27^, and the PD-associated *LRRK2* variants can either associate with a synucleinopathy or a pure tauopathy with the accumulation of AD-type tau^28^. Pathologic forms of ɑS, Aβ, and tau synergistically interact with one another, enhancing aggregation and neurotoxicity and suggesting the presence of a shared mechanistic link among proteinopathies^29,30^. Biophysical data suggesting that aggregation-prone proteins can cross-fibrillize – for example, Aβ with ɑS^31^ and ɑS with tau^32^-indicates a possible mechanism to explain the coexistence of proteinopathies. How closely these *in vitro* studies mimic the physiologic interactions *in vivo* remains unclear, and our understanding of the shared genetic basis of different synucleinopathies and across proteinopathies remains far from complete.

The field has employed two distinct but complementary genetic approaches to understand the complex mechanisms driving neurodegenerative diseases: human genetic analysis and genetic analysis of tractable model systems. The most comprehensive modifier screens to date in model organisms have been performed in yeast in which the expression of ɑS and Aβ in yeast leads to robust cytotoxicity^33^. Genes encompassing ∼85% of the yeast proteome have been individually co-expressed with each of these toxicity proteins to identify genetic modifiers of ɑS^34^ and Aβ^35^ toxicity, respectively. The system offers a convenient way to isolate the cytotoxicity of ɑS from Aβ because these proteins do not natively exist in this cell. Strikingly, despite this absence, the system is disease-relevant because the Aβ screen^34,36,37^ statistically enriched for genes implicated in AD (*CD2AP, PICALM, INPP5D, RIN3*) and the ɑS screen for PD/parkinsonism-implicated genes^35,36^ (*ATP13A2, CHCHD2, SYNJ1, VPS35*^35,37^). Interestingly, in these acute cytotoxicity models, there was little meaningful overlap between ɑS and Aβ screen hits. Notable exception was a modification of both ɑS and Aβ proteotoxicity by paralogs, *ROM1* and *ROM2*^38^ guanidine exchange factors of the yeast Rho GTPase Rho1p involved in actin cytoskeleton regulation^36^.

In human genetics, the standard method has been genome-wide association studies (GWAS). These are best powered for common variants (with MAF >1%). However, such variants typically have small effect sizes and are located in non-coding genomic regions, thus making it challenging to map them back to specific genes. To identify rare variants, very large cohorts comprising of whole-genome or whole-exome sequence data are required for late-onset complex diseases. Recent investigations have attempted to narrow the search space for rare variants to circumvent this. For example, variants have been restricted to particular classes of variants^39^, confined to restricted ontologies emerging from common-variant GWAS^40^, and anchored to specific cellular processes^41^or organelles. One such study^42^ targeted exome sequencing of lysosomal genes demonstrated the importance of this organelle to PD, a finding subsequently verified more definitively in larger meta-GWAS analysis^43^.

To examine whether the ɑS and Aβ proteotoxicity modifiers and gene networks can yield important insights into shared risk across synucleinopathies and between PD and AD, we performed targeted exome screening of human orthologs of these modifier genes as well as additional known AD, PD, and neurodegeneration-related genes. We targeted a relatively small but deeply characterized cohort of 496 patients with different synucleinopathies (PD, DLB, and MSA). We performed carefully controlled joint calling with ∼2,500 aged neurotypical controls from the Medical Genome Reference Bank (MGRB)^44^. We selected a wide range of orthologs of the yeast genes to capture broad expression across tissues and CNS cells because we hypothesized that distinct cell types would be involved in different synucleinopathies. Our goal was to reduce the sequencing sampling space to a set of genes known *a priori* to be enriched with known PD and AD Mendelian genetic risk factors to enable us to find meaningful signals even in a modestly sized cohort. Critically, we matched proteinopathy screening in yeast to a molecular, rather than clinical, phenotype in patients. Put another way, we matched synucleinopathy in humans, regardless of clinical diagnosis, to screens specifically against proteinopathies in yeast.

Reassuringly, our cross-species screen identified significant enrichment of rare nonsynonymous variants in known PD genes *GBA1* and *LRRK2*. However, three AD genes (*CD33, CR1,* and *PSEN2*) were also implicated, building on emerging genetic evidence for shared risk across AD and PD. Our statistically significant genes crossed over synucleinpathy boundaries and implicated shared genetic risk among PD, DLB and MSA. Most strikingly, when we validated our findings in orthogonal (UK biobank and AMP-PD) cohorts, our statistical signal was stronger for Aβ toxicity modifiers than ɑS. These modifiers centered on genes that, together with *PSEN2*, have been strongly implicated in the regulation of actin cytoskeleton (*ARGHEF1, ARHGEF28, PKN2, PRKCD, PASK*), genes that collectively we refer to as the *PSEN2*/Aβ-actin gene module. Three of these genes were positive regulators or effectors of RhoA, a central actin cytoskeleton signaling factor. Patients with mutations in these genes have diffuse synucleinopathy with brainstem and cortical pathology and in human neurons, simply downregulating RhoA can deplete stabilized F-actin and enhance ɑS pathology. Among this gene module, *PSEN2* itself emerged as the clearest risk factor, essential for cortical and dopaminergic (DA) neuron survival, downregulated in cortical and dopaminergic neurons with contributions to aggregated PD GWAS risk and, finally, tied to regulation of ɑS expression itself.

## RESULTS

To investigate potentially shared genetic risk and penetrance factors for the synucleinopathies of interest, we combined functional genomics with human genetics. We leveraged the largely non-overlapping modifiers from genome-wide over-expression screens of ɑS^35^ and Aβ^34^ proteotoxicity in yeast cells to probe the proteotoxicities of interest, rather than be limited to clinical diagnosis. Since yeast cells lack clear homologs of Aβ and ɑS, we sequenced a wide range of human orthologs for these genes, including the known Mendelian genetic risk factors for AD and PD among them^36^. This set was further enhanced by including known AD, PD, and ataxia risk genes. We performed high-depth sequencing (∼100x) of a total of 430 proteotoxicity modifier genes in a human genetic screen of 496 patients with synucleinopathies (PD, DLB and MSA) (Figure 1A). We jointly-called these patients with 2,516 aged (>70 years old) unrelated individuals with no history of cancer, cardiovascular disease, or dementia from the Medical Genome Reference Bank (MGRB)^44^ by extracting the same target regions from their whole genomes (Figure 1A, Methods). Samples from HapMap with northern and western European ancestry (CEU) samples were also included for quality control purposes. We hypothesized that reducing the sequencing sampling space to a set of genes known *a priori* to be enriched with known PD and AD Mendelian genetic risk factors would enable us to find meaningful signals even in a modestly sized cohort.

**Figure 1.**
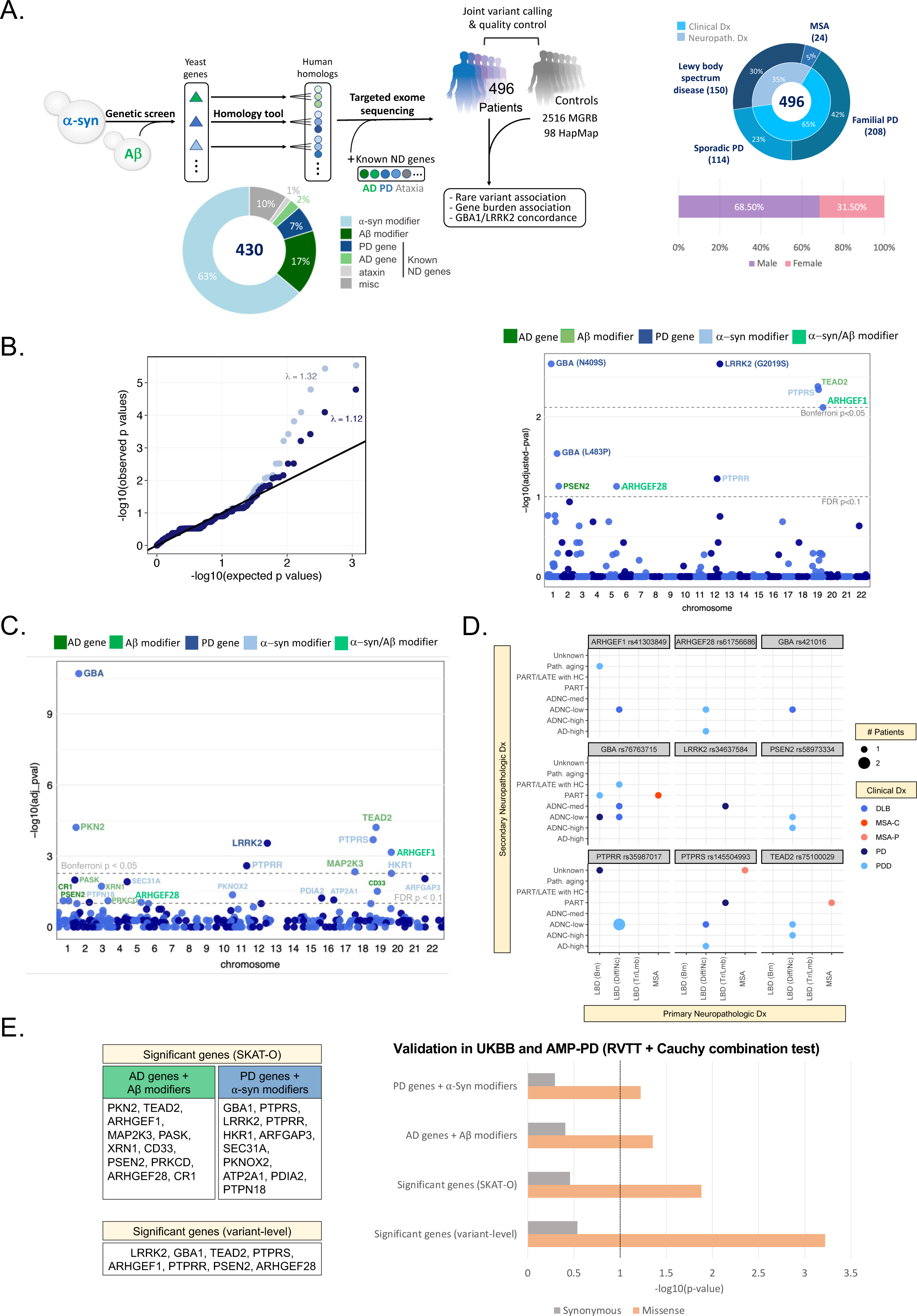
High-depth (∼100X) patient exome screening identifies shared genetic risk among synucleinopathies, and an overlap with known PD and AD genes. (A) Study design (sequencing and analysis workflow), breakdown of target genes, and primary diagnosis and demographics of the sequenced synucleinopathy cases. (B) Quantile-quantile plot of observed versus expected p-values from the variant-level Fisher’s Exact test for association between cases and MGRB controls and their respective genomic inflation with and without ClinVar variants (left). Manhattan plot of FDR-adjusted (permutation-based case-control resampling) p-values resulting from the variant-level association (right). Dashed lines indicate Bonferroni and FDR significance thresholds. Amino acid changes are noted in parentheses for known ClinVar pathogenic variants. Eight missense variants and one splice region variant were significant at FDR adjusted p-value < 0.1. (C) Manhattan plot showing FDR-adjusted p-values from the gene-level rare variant association test (cases vs. MGRB controls) using SKAT-O. Dashed lines indicate Bonferroni and FDR significance thresholds. At FDR adjusted p-value < 0.1, 22 significant genes were identified to have increases rare nonsynonymous variant burden in cases. (D) Top variant-level hits are shared across clinical synucleinopathy diagnoses in pathologically confirmed cases. Each dot is colored by the clinical diagnosis. Size of each dot indicates the number of patients with a particular diagnosis. Primary neuropathological diagnoses are shown on the x-axis and secondary neuropathological diagnoses are shown on the y-axis. See Supplemental Figure 1 for the expanded list. Here, ADNC: Alzheimer disease neuropathologic change; LATE: limbic predominant age-related TDP-43 encephalopathy; MCI: mild cognitive impairment; MSA-C: multiple system atrophy cerebellar type; MSA-P: multiple system atrophy parkinsonian type; PART: primary age-related tauopathy (tau deposits found without beta-amyloid); PDD: Parkinson’s disease with dementia, LBD (Brn): LBD (Brainstem), LBD (Diff/Nc): LBD (Diffused/Neocortical), and LBD (Tr/Lmb): LBD (Translational/Limbic), Path. aging: pathologic aging with beta-amyloid deposits but no tau. (E) Validation of variant- and gene-level rare variant burden in Parkinson’s disease (PD) in the UK Biobank (UKBB) and AMP-PD datasets using the rare variant trend test (RVTT). We tested variant-level significant-genes, SKAT-O significant genes, as well as two subsets of significant genes: (i) AD genes and Aβ modifiers, and (ii) PD genes and α-syn modifiers. We observed a significant increasing trend in rare missense variant occurrences in PD cases compared to control in both cohorts. There was no enrichment of synonymous variants. Cauchy combination test was performed to combine the RVTT p-values from individual datasets. Combined p-values on a negative log10 scale are shown in the bar chart.

### Increased rare variant burden in both PD and AD genes and in both Aβ and ɑS toxicity modifiers across synucleinopathies

To assess the contribution of the Aβ and ɑS genetic networks to the risk of disease in synucleinopathies and identify shared genetic drivers across different synucleinopathies, we performed the one-sided Fisher exact test on each biallelic variant with a minor allele frequency (MAF) cut-off of 0.01 both at single-variant and gene levels on the European cases vs controls (Ncase = 496 and Ncontrol = 2,516). Singleton variants i.e., variants that appeared exactly once in both PD cases and MGRB controls were excluded because of the significant deviation observed in their distribution between cases and controls (see Methods).

We identified several screen-wide significant variants (FDR-adjusted p-value < 0.1) in patients compared to controls (Figure 1B). Unsurprisingly, the top variants were the known PD-associated missense variants --- *LRRK2* rs34637584 (G2019S), *GBA1* rs76763715 (N409S) and rs421016 (L483P). Excluding the variants of clinical importance (i.e., variants reported as “pathogenic” in ClinVar), we identified several significant missense and splice variants in Aβ modifiers/AD genes: *TEAD2, PSEN2*; ɑS modifiers/PD genes: *GBA1*, *PTPRR, PTPRS*; and also *ARHGEF1, ARHGEF28* which are modifiers of both Aβ and ɑS toxicity (Figure 1B, right; Table 1). The exclusion of ClinVar variants also helped us reduce the inflation caused by including AD and PD-related genes in the target gene set (λ with ClinVar variants: 1.324, λ without ClinVar variants =1.123; Figure 1B, left), suggesting that the residual inflation is in fact arising from true enrichment of causal variants.

**Table 1.** Significant rare nonsynonymous variants (MAF < 0.01) in synucleinopathy cases versus controls. Fisher exact test was performed to test the association of variants with disease risk. Significant variants were selected with a cutoff of FDR-adjusted p-value < 0.1.

| CHR:POS:REF:ALT | rsid | AA<br>change | CSQ | Gene | pval | FDR-adj<br>pval | Description | Rationale for<br>sequencing |
| --- | --- | --- | --- | --- | --- | --- | --- | --- |
| 12:40734202:G:A | rs34637584 | G2019S* | Missense | LRRK2 | 3.30x10<sup>-06</sup> | 2.17x10<sup>-03</sup> | Vesicle trafficking | PD risk gene |
| 1:155205634:T:C | rs76763715 | N409S* | Missense | GBA1 | 4.27x10<sup>-06</sup> | 2.17x10<sup>-03</sup> | Lysosomal hydrolase | PD risk gene |
| 19:49852030:C:G | rs75100029 | G226A | Missense | TEAD2 | 1.23x10<sup>-05</sup> | 4.18x10<sup>-03</sup> | Transcription factor,<br>Hippo signaling | Homolog of Aβ<br>modifier *Tec1*<br>(OE=Supp) |
| 19:5240271:G:A | rs145504993 | P549L | Missense | PTPRS | 1.79x10<sup>-05</sup> | 4.56x10<sup>-03</sup> | Receptor tyrosine<br>phosphatase, MAPK<br>signaling | Homolog of α-syn<br>modifier *Ptp2*<br>(OE=Supp) |
| 19:42392819:C:G | rs41303849 |  | Splice<br>region<br>variant | ARHGEF1 | 3.74x10<sup>-05</sup> | 7.63x10<sup>-03</sup> | RhoA / actin<br>cytoskeleton<br>regulation | Homolog of Aβ<br>modifier *Rom1*<br>(OE=Enh) and α-syn<br>modifier *Rom2*<br>(OE=Enh) |
| 1:155205043:A:G | rs421016 | L483P* | Missense | GBA1 | 1.70x10<sup>-04</sup> | 2.88x10<sup>-02</sup> | Lysosomal hydrolase | PD risk gene |
| 12:71139860:A:G | rs35987017 | Y249H | Missense | PTPRR | 4.08x10<sup>-04</sup> | 5.94x10<sup>-02</sup> | Receptor tyrosine<br>phosphatase | Homolog of α-syn<br>modifier *Ptp2*<br>(OE=Supp) |
| 1:227071449:G:A | rs58973334 | R62H | Missense | PSEN2 | 5.80x10<sup>-04</sup> | 7.39x10<sup>-02</sup> | Gamma secretase,<br>notch/APP<br>processing, actin<br>cytoskeleton<br>regulation | AD risk gene |
| 5:73205247:C:A | rs61756686 | T1391N | Missense | ARHGEF28 | 6.55x10<sup>-04</sup> | 7.42x10<sup>-02</sup> | RhoA / actin<br>cytoskeleton<br>regulation | Homolog of Aβ<br>modifier *Rom1*<br>(OE=Enh) and α-syn<br>modifier *Rom2*<br>(OE=Enh) |

\* ClinVar “pathogenic” variants

**Table 2.** Significant genes with rare nonsynonymous variant burden in synucleinopathy cases versus controls. Missense, nonsense, and splice variants with MAF < 0.01 were collapsed and gene-level SKAT-O test was performed, and significant genes were selected with a cutoff of FDR-adjusted p-value < 0.1.

| Gene | pval | Bonferroni<br>pval | FDR-adj<br>pval | Description | Rationale for sequencing |
| --- | --- | --- | --- | --- | --- |
| GBA1 | 6.67x10<sup>-14</sup> | 1.96x10<sup>-11</sup> | 1.96x10<sup>-11</sup> | Lysosomal hydrolase | PD risk gene |
| PKN2 | 4.70x10<sup>-07</sup> | 1.38x10<sup>-04</sup> | 6.18x10<sup>-05</sup> | Ser/Thr-protein kinase, Rho/Rac effector protein, **RhoA / actin cytoskeleton regulation** | Homolog of Aβ modifier *Pkc1* (OE=Enh) |
| TEAD2 | 6.32x10<sup>-07</sup> | 1.85x10<sup>-04</sup> | 6.18x10<sup>-05</sup> | Transcription factor, Hippo signaling | Homolog of Aβ modifier *Tec1* (OE=Supp) |
| PTPRS | 2.80x10<sup>-06</sup> | 8.20x10<sup>-04</sup> | 2.05x10<sup>-04</sup> | Receptor tyrosine phosphatase, MAPK signaling | Homolog of α-syn modifier *Ptp2* (OE=Supp) |
| LRRK2 | 4.91x10<sup>-06</sup> | 1.44x10<sup>-03</sup> | 2.88x10<sup>-04</sup> | Vesicle trafficking | PD risk gene |
| ARHGEF1 | 1.42x10<sup>-05</sup> | 4.16x10<sup>-03</sup> | 6.93x10<sup>-04</sup> | **RhoA / actin cytoskeleton regulation** | Homolog of Aβ modifier *Rom1* (OE=Enh) and α-syn modifier *Rom2* (OE=Enh) |
| PTPRR | 6.17x10<sup>-05</sup> | 1.81x10<sup>-02</sup> | 2.58x10<sup>-03</sup> | Receptor tyrosine phosphatase | Homolog of α-syn modifier *Ptp2* (OE=Supp) |
| MAP2K3 | 1.27x10<sup>-04</sup> | 3.72x10<sup>-02</sup> | 4.64x10<sup>-03</sup> | Dual specificity kinase, MAPK signaling | Homolog of Aβ modifier *PBS2* (OE=Enh) |
| HKR1 | 1.67x10<sup>-04</sup> | 4.88x10<sup>-02</sup> | 5.43x10<sup>-03</sup> | Zinc finger protein | Homolog of α-syn modifier *Tda9* (OE=Supp) |
| ARFGAP3 | 3.10x10<sup>-04</sup> | 9.09x10<sup>-02</sup> | 9.09x10<sup>-03</sup> | GTPase activator, Golgi vesicle-mediated transport | Homolog of α-syn modifier *Glo3* (OE=Enh) |
| PASK | 3.89x10<sup>-04</sup> | 0.113902 | 1.04x10<sup>-02</sup> | Ser/Thr-protein kinase, energy homeostasis | Homolog of Aβ modifier *Psk1* (OE=Enh) |
| SEC31A | 5.01x10<sup>-04</sup> | 0.146700 | 1.22x10<sup>-02</sup> | Component of coat protein complex II, ER-Golgi transport vesicles | Homolog of α-syn modifier *Sec31* (OE=Enh) |
| XRN1 | 8.45x10<sup>-04</sup> | 0.247499 | 1.90x10<sup>-02</sup> | Exonuclease, mRNA metabolism | Homolog of Aβ modifier *Xrn1* (OE=Enh) |
| CD33 | 1.48x10<sup>-03</sup> | 0.434354 | 3.10x10<sup>-02</sup> | SIGLEC family, innate immune system | AD risk gene |
| PKNOX2 | 2.24x10<sup>-03</sup> | 0.655811 | 4.37x10<sup>-02</sup> | Sequence-specific DNA binding, actin monomer binding, cell proliferation, differentiation, and death | Homolog of α-syn modifier *Cup9* (OE=Supp) |
| ATP2A1 | 3.28x10<sup>-03</sup> | 0.960284 | 6.00x10<sup>-02</sup> | Calcium ion binding, nucleotide binding | Homolog of α-syn modifier *Pmr1* (OE=Enh) |
| PDIA2 | 4.19x10<sup>-03</sup> | 1 | 7.23x10<sup>-02</sup> | ER protein, catalyzes protein folding | Homolog of α-syn modifier *Epd1* (OE=Enh) |
| PSEN2 | 5.12x10<sup>-03</sup> | 1 | 7.78x10<sup>-02</sup> | Gamma-secretase complex, cleaves Ab precursor protein, **actin cytoskeleton regulation** | AD risk gene |
| CR1 | 5.29x10<sup>-03</sup> | 1 | 7.78x10<sup>-02</sup> | Complement cascade, innate immune system | AD risk gene |
| PRKCD | 5.31x10<sup>-03</sup> | 1 | 7.78x10<sup>-02</sup> | Ser/Thr-protein kinase, apoptosis, **actin cytoskeleton regulation** | Homolog of Aβ modifier *Pkc1* (OE=Enh) |
| ARHGEF28 | 6.58x10<sup>-03</sup> | 1 | 9.07x10<sup>-02</sup> | **RhoA / actin cytoskeleton regulation** | Homolog of Aβ modifier *Rom1* (OE=Enh) and α-syn modifier *Rom2* (OE=Enh) |
| PTPN18 | 6.81x10<sup>-03</sup> | 1 | 9.07x10<sup>-02</sup> | Protein tyrosine phosphatase, signaling | Homolog of α-syn modifier *Ptp2* (OE=Supp) |

To identify gene-level rare variant burden in synucleinopathies, we performed the sequence kernel association test-optimized (SKAT-O) test by collapsing the nonsynonymous variants (missense, nonsense, and splice) within each gene (see Methods). Synonymous variants were tested as an internal control. We identified screen-wide significant associations between the synucleinopathy risk and the burden of nonsynonymous variants in 22 genes (FDR-adjusted p-value < 0.1, Figure 1C): two were known PD genes (*GBA1*, *LRRK2*), three were surprisingly known AD genes (*CR1*, *CD33*, *PSEN2*), nine were modifiers of ɑS toxicity (*PTPRS, PTPRR, HKR1, ARFGAP3, SEC31A, PKNOX2, PDIA2, ATP2A1, PTPN18*), six were modifiers of Aβ toxicity (*TEAD2, PKN2, MAP2K3, PASK, XRN18, PRKCD*) and two were modifiers of both ɑS and Aβb toxicity (*ARHGEF1* and *ARHGEF28*). Nine of these genes --- *GBA1, PKN2, TEAD2, PTPRS, LRRK2, ARHGEF1, PTPRR, MAP2K3, HKR1* --- were Bonferroni significant (Bonferroni-p < 0.05). There was no significant enrichment of synonymous variants in synucleinopathy cases in these genes. None of these genes, except for *GBA1* and *LRRK2*, has been identified in large-scale rare-variant studies of PD^45,46^ (Supplementary Table 1).

### Genes with significant rare variant burden cross clinicopathologic disease boundaries

Using 151 of our cases that had additional neuropathologic data, we asked whether the significant variants we identified, crossed clinical disease boundaries. In pathologically confirmed cases, these significant variants indeed crossed different synucleinopathies (Figure 1D). Additionally, among the 11 patients with top variants in either AD genes (*PSEN2*) or in the Aβ-modifier genes (*TEAD2, ARHGEF1, ARHGEF28*), only three had concomitant AD pathology (Figure 1D, Supplementary Figure 1; see also Figure 4).

As cases with known pathogenic mutations may harbor additional risk variants, we also tested whether we could find additional variants associated with familial forms of synucleinopathies with known *GBA1* and *LRRK2* variants. Penetrance of the PD phenotype in carriers of these mutations is highly variable (10-50%)^47–49^. The G*BA1* N409S and *GBA1* L483P mutations were detected in 13/496 (2.6%) and 9/496 (1.8%) of our cases compared to 10/2516 (0.39%) and 6/2516 (0.23%) of MGRB controls, respectively (Supplementary Figure 2). Of the 21 cases (one case harbored both the *LRRK2* G2019S and *GBA1* L483P variant), 8 were sporadic PD, 7 LBD, 6 familial PD, and 1 MSA. The *LRRK2 G2019S* was present in a total of 9/496 (1.8%) cases, all familial PD, with the G2019S mutation and 3/2516 (0.12%) controls (Supplementary Figure 2).

Comparing all 22 cases with *GBA1* mutations with all other cases as well as GBA1-N409S and *GBA1*-L483P carriers separately using Cohort Allelic Sums Test (CAST)^50^, we identified two missense variants to be significantly enriched (Bonferroni adjusted p<0.05) in *GBA1*-N409S carriers compared to all other cases: *PTPRB* (rs202134984), and *FREM2* (rs150928081). Additionally, performing gene-level burden tests, we identified one additional significant hit, *MICAL3* and two additional nominally significant genes: *PTPRB* and *PRKCD* (Supplementary Table 2). *GBA1*-N409S carriers drove the association, as these genes were significantly enriched in N409S cases, and no genes were significant in *GBA1*-L483P carriers versus all other cases. Performing CAST in *LRRK2* G2019S carriers versus all other cases, we observed nominally significant burden of rare nonsynonymous SNVs (missense, splice, stop gained, stop lost, start lost), MAF<1% in *PICALM* (rs34013602), *PKN2* (rs200490316) and *ATP12A* (rs61998252, rs61740542). No additional genes were identified in gene-level burden testing.

### Rare variant trend test validates significant both ɑS and Aβ gene modules in UK Biobank and AMP-PD

We validated our variant- and gene-level significant results in two independent PD datasets: UK Biobank (UKBB) ∼500K whole exomes and Accelerating Medicines Partnership for PD (AMP-PD) whole genomes. Large-scale independent studies of these datasets^45,46^ found no significant genes at single gene level except *GBA1* and *LRRK2* (Supplementary Table 1). Thus, we performed the pathway-based rare variant trend test (RVTT)^51,52^ with the genes identified by variant and gene-level tests in our original cohort. RVTT selects qualifying rare variants using a variable minor allele frequency (MAF) threshold approach and uses the Cochran-Armitage test statistic to measure a group’s increased burden of variants. RVTT provides permutation-based p-values as an indicator of the strength of association.

In the UKBB dataset, we compared 2,273 unrelated European PD cases (ICD10 code: G20) of age 40 or older with 6,711 controls randomly sampled from 167,188 unrelated European neurotypical (no ICD10 code between G01-G99) individuals of age 60 or older (Methods). From the AMP-PD dataset, we compared 1,598 unrelated European sporadic PD cases with 1,095 age-matched neurotypical controls. HBS samples were excluded from the analysis due to potential overlap with our original sequencing cohort (Methods).

We applied RVTT on each dataset individually and meta-analyzed the p-values with the Cauchy combination test (CCT)^53^. We observed a significant enrichment of rare missense variants (MAF < 0.01) in the significant genes identified at both variant- and gene-levels in the combined analysis (Figure 1E). To assess the contribution of significant genes, other than *LRRK2* and *GBA1*, to disease risk, we also ran RVTT on two subsets of significant genes (SKAT-O): (i) AD genes and Aβ-modifiers and (ii) PD genes and ɑS-only modifiers. Both were significant at CCT p-value cutoff of 0.1. Surprisingly, the set of known AD genes and Aβ-modifiers showed a stronger association to PD risk compared to the known PD genes and ɑS-only modifiers (Figure 1E). Detailed results from both cohorts are available in Supplementary Table 3. No significant trend was observed in synonymous variants between cases and controls in the gene sets under question. This provides evidence for the association of our significant AD-related and Aβ-modifying genes with the risk of PD. Notably, six of these genes, namely *ARHGEF1, ARHGEF28, PKN2, PASK, PRKCD* and *PSEN2*, have been tied to the regulation of actin cytoskeleton (Figure 2).

**Figure 2.**
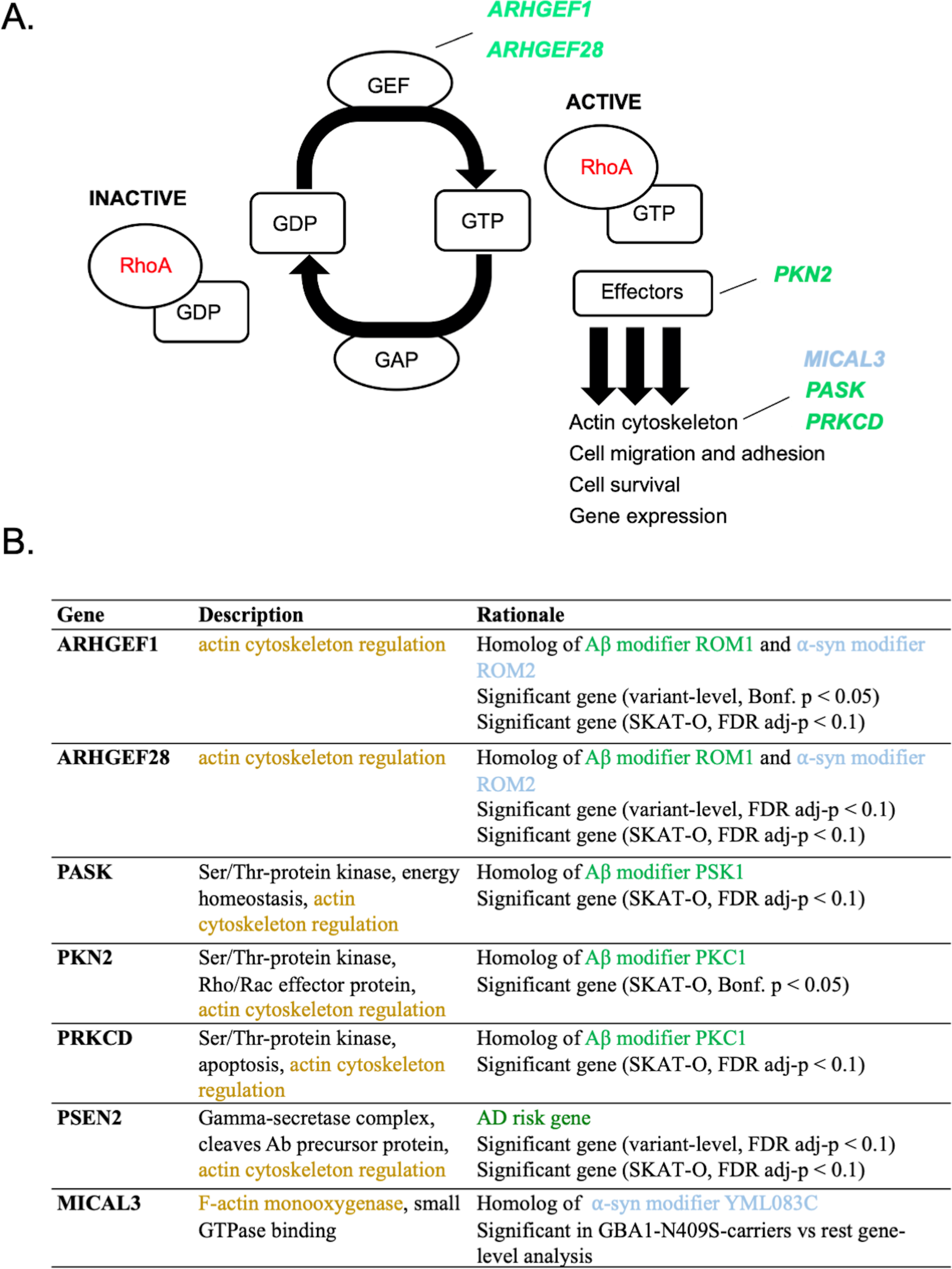
A gene module comprising RhoA signaling/actin cytoskeleton regulators previously tied to AD and A**β** toxicity are among the more significant genes identified in this study. (A) Schematic diagram of Ras homology family member A (RhoA) signaling. RhoA is a master regulator of actin cytoskeleton stabilization and other processes. Signaling is activated by guanidine exchange factor Arhgef proteins. A key effector of actin cytoskeleton changes is the Pkn2 kinase. (B) Genes with significant rare variant burden in our analysis, including the known AD gene *PSEN2,* that have been mechanistically connected to regulation of the actin cytoskeleton.

### Common regulatory changes near significant Aβ/AD genes

Recently, there is growing evidence that, even for rare-variants, impact on human disease ultimately leads to altered gene expression^54,55^. However, with existing sample sizes and statistical tools, we are not powered to detect the regulatory roles of rare variants directly. Common regulatory variations such as expression quantitative trait loci (eQTLs) and splice quantitative trait loci (sQTLs) can provide a way to connect genes enriched in rare variants with expression changes in disease-relevant tissues and cell types. Hence, we asked whether the genes recovered in our analysis were enriched for such common regulatory variations. We focused on known AD genes and Aβ toxicity modifiers because this was where we saw the most convincing signal in our UKBB/AMP-PD analysis (Figure 1E). We interrogated the GTEx Brain and PsychENCODE datasets and identified a significant enrichment (*p*=5.18×10^-3^, Supplementary Figure 3A), indicating a propensity of tissue-relevant genetic regulation control of these significant genes from our rare-variant analysis. Furthermore, we observed many of these genes have significant cell-type-specific cis-eQTLs in different brain cell types (Supplementary Figure 3B, Suppl. Table 4) in two publicly available datasets of single-cell eQTLs from prefrontal cortex^56^ and mid-brain^57^. In the mid-brain dataset, filtering for PD GWAS variants overlapping with single cell eQTLs near our known AD genes and Aβ toxicity modifiers, we detected significant enrichment with loci at different PD GWAS p-value thresholds < 1×10^-5^ (Supplementary Figure 3C) in dopaminergic neurons (DAs), suggesting a relationship between PD risk modifying regulatory variants acting in the DA neuron context through the AD and Aβ toxicity-modifier genes. Notably, this enrichment was primarily driven by *PSEN2* which also had multiple significant cis-eQTLs in excitatory glutamatergic neurons (Supplementary Table 4).

### Shared synucleinopathy risk factors comprise a *PSEN2*/Aβ-Actin regulator module that includes RhoA regulators and effectors

Our initial screen and RVTT validation, together with eQTL analysis, pointed to AD and Aβ modifier genes as being the strongest contributor to synucleinopathy risk among our targeted genes. This was an altogether unexpected finding. When we perused the list more closely, we were struck by one common thread trying most of these genes together, namely a link to regulation of the actin cytoskeleton (see Figure 2). Dysregulation of the cytoskeleton has been linked repeatedly to neurodegeneration and specifically to synucleinopathy in in vitro, cellular, and in vivo models. Most intriguing to us, three of the genes we identified encoded proteins Arhgef1/28 and Pkn2 that were regulators and effectors of RhoA signaling, respectively. Psen2 was selected to analyze along with these genes because it is a known AD risk factor, but because it also has been shown to directly interact with the filamin class of actin-binding proteins. We thus established a human iPSC-based synucleinopathy model to further investigate these relationships.

### A tractable synucleinopathy hiPSC model is established with transgenic *SNCA* expression

iPSC modeling for neurodegenerative proteinopathies suffers from poor reproducibility and tractability, and low levels of endogenous αS. Recently, we^58^ and others^59,60^ have begun to establish a suite of tractable models to study different aspects of neurodegenerative disease biology. We extended this development by engineering the WTC11 iPSC line that was selected for engineering by the Allen Institute for Cell Science (https://www.allencell.org/cell-catalog.html). It is known that simply inheriting extra copy number variants (CNVs) of wild-type αS through gene-multiplication of the *SNCA* locus can lead to PD, PDD or DLB. These patients even have MSA-like glial cytoplasmic pathologies^61^. To create a better surrogate for this pathobiology, we established Cortical induced synucleinopathy (CiS) models (Figure 3A). Cortical glutamatergic neurons are straightforward to generate through one-step trans-differentiation with Ngn2 and exhibit αS pathology across synucleinopathies, thus providing a good substrate for rapid modeling of synucleinopathy “in the dish”.

**Figure 3.**
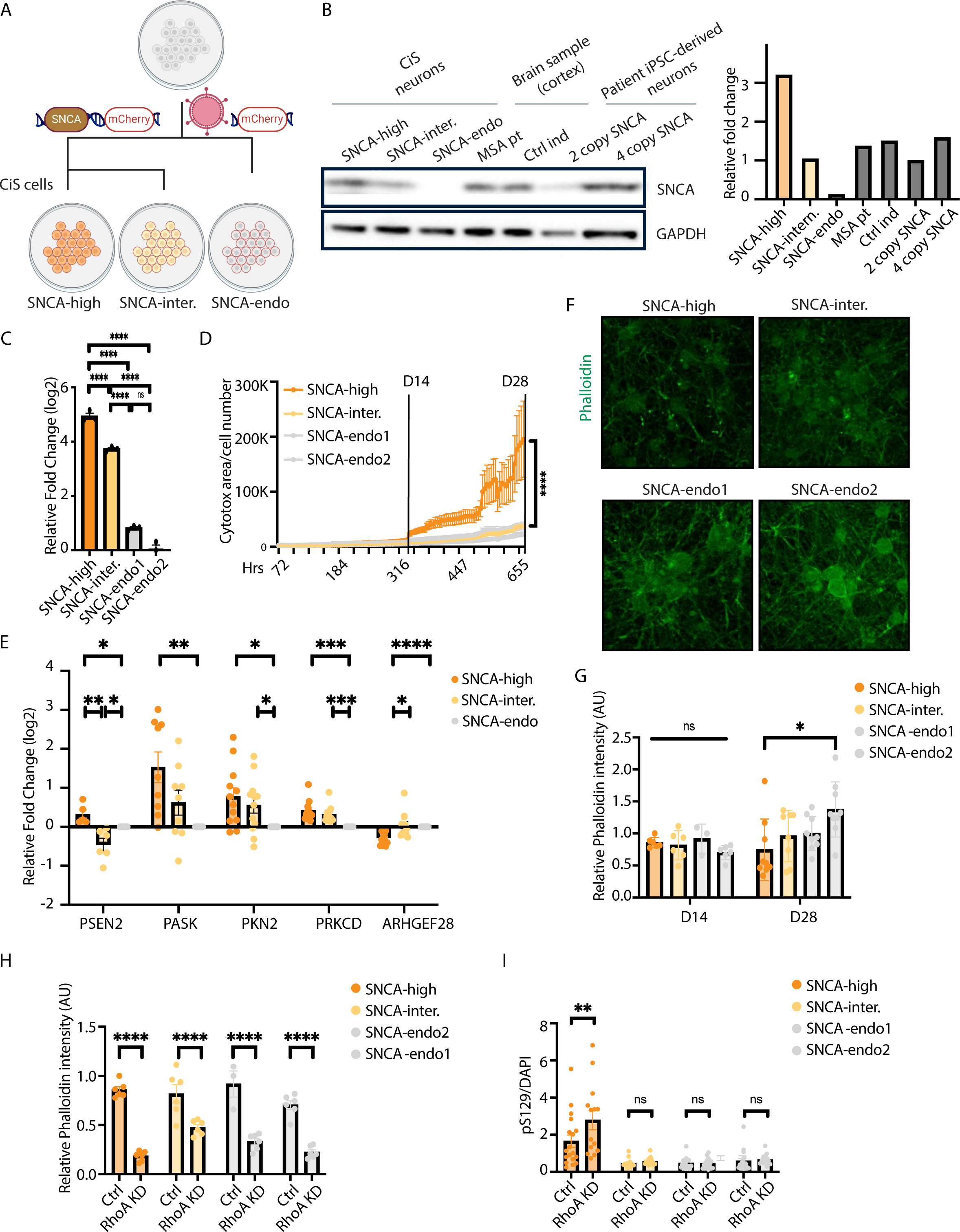
RhoA downregulation phenocopies *SNCA*-overexpression and induces pathologic ɑS accumulation in human neurons. (A) Schematic diagram: constructing a tractable **C**ortical neuron **i**nduced **S**ynucleinopathy (CiS) model through viral induction of *SNCA* or control gene (together with IRES mCherry) into a widely used parental (WTC11) iPSC. This line harbors Ngn2 at a safe-harbor *AAVS1* locus enabling rapid induction of glutamatergic neurons upon transient exposure to doxycycline. (B) Whole-cell western blot shows comparable αS level in CiS neurons to human cortex samples and to iPSC-derived neurons from an *SNCA* triplication carrier. (C) Relative mRNA expression level of *SNCA* in CiS neurons. Values represent mean ± s.e.m. (n = 3; ****p < 0.0001; one-way ANOVA). (D) Cytotox assay of DIV1-DIV28 CiS neurons shows significantly increased cellular death in *SNCA*-high neurons starting from DIV14. Values represent mean ± s.e.m. (****p < 0.0001; two-way ANOVA). (E) Relative mRNA expression level of the identified *PSEN2*/Aβ-actin gene module in D28 CiS neurons. Values represent mean ± s.e.m. (n = 3; *p < 0.05; **p < 0.01; ***p < 0.001; ****p < 0.0001; t-test). (F) Sample confocal images of DIV28 CiS neurons immunostained with Phalloidin. (G) Immunostaining quantification analysis shows at DIV28, Phalloidin intensity of *SNCA*-high neurons is significantly lower than that of *SNCA*-endo neurons. However, there is no difference detected in DIV14 neurons. Values represent mean ± s.e.m. (*p < 0.05; one-way ANOVA). (H-I) Immunostaining quantification shows RhoA knockdown (H) significantly decreased Phalloidin intensity in all DIV14 CiS neurons (phenocopying the effect of *SNCA* overexpression); (I) selectively increased pS129+ cells in DIV14 *SNCA*-high neurons. Values represent mean ± s.e.m. (*p < 0.05; **p < 0.01; ****p < 0.0001; two-way ANOVA).

**Figure 4.**
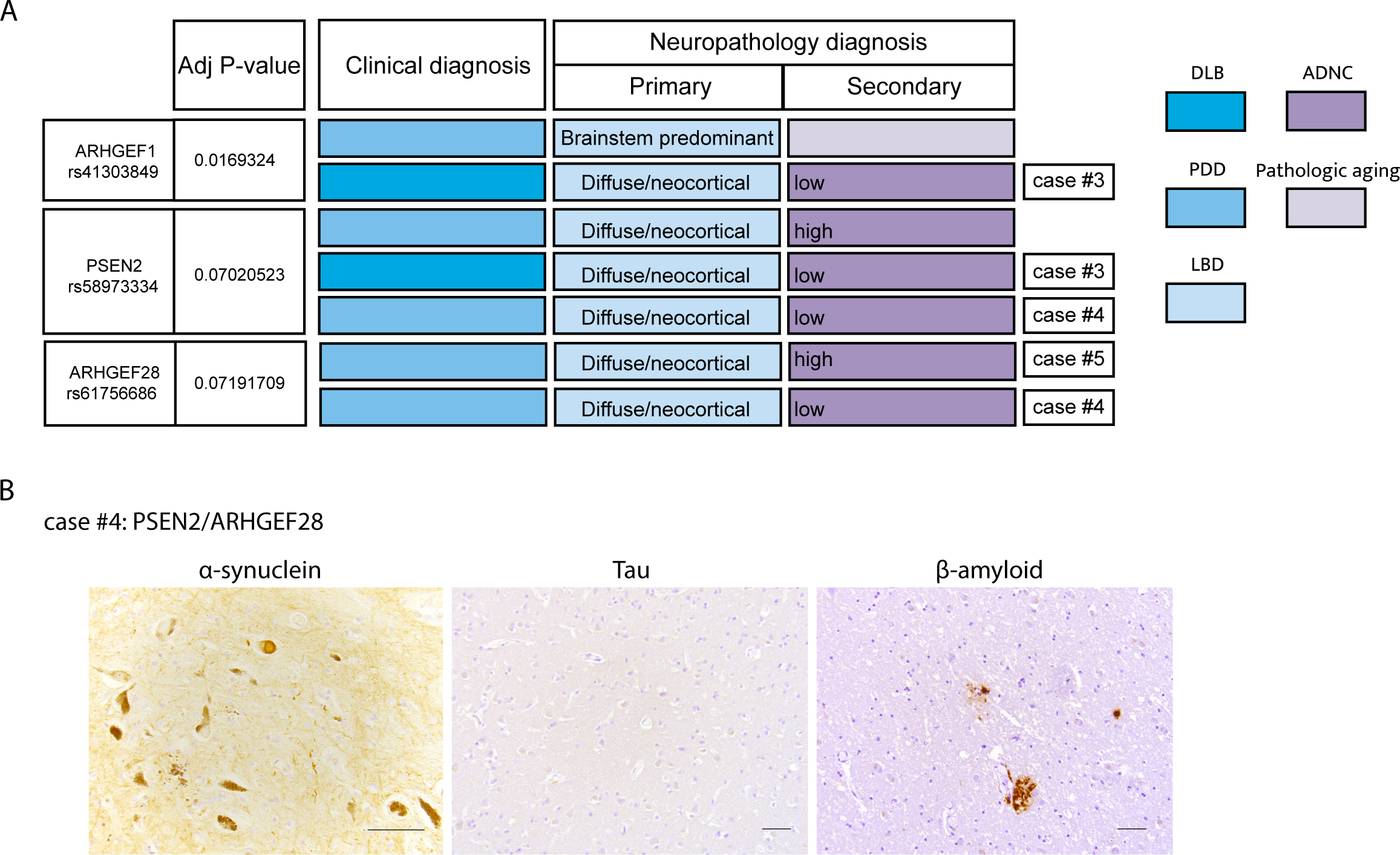
Clinical and neuropathology diagnosis of patients carrying *ARHGEF1/28*, *PSEN2* variants. (A) The distinction between PD with dementia (PDD) and DLB is based on clinical criteria. Neuropathologic diagnosis for Lewy Body Disease (LBD) is either present/absent, and the type of LBD (brainstem predominant, limbic, amygdala only, or diffuse/neocortical). Neuropathologic criteria for Alzheimer’s disease (AD) outlines three “scores” corresponding to Amyloid/Thal, Braak/tau and CERAD/neuritic plaque. Based on these scores, cases are designated as having no, low, intermediate, or high level of AD neuropathologic change (ADNC). No/low ADNC is generally associated with cognitively normal individuals. Pathologic aging: beta-amyloid deposits being found without any tau. Lower panel: Histopathology from *PSEN2* variant carrier, which is an example of LBD and low level ADNC. (B) Histopathology from case #4, a carrier of *PSEN2* and *ARHGEF28* variants. Low ADNC was observed in this case (scalebar: 50 um).

WTC11 hiPSCs with doxycycline-inducible-*NGN2* knocked-in at the *AAVS1* safe-harbor locus were transduced with a lentiviral *SNCA-IRES-mCherry* transgenic construct or with *mCherry* alone serving as a control. hiPSCs were flow cytometry-sorted utilizing the mCherry fluorescence and replated at low density for single-cell clonal selection. iPSC clones were expanded, karyotyped, and examined for αS expression. Two iPSC clones in *SNCA*-overexpression group, two clones in control group were selected for further investigation and named *SNCA-high, SNCA-intermediate, SNCA-endogenous1, SNCA-endogenous2*, accordingly.

We compared αS expression levels in DIV28 neurons with human brain samples and with 4-copy *SNCA* neurons from a patient harboring *SNCA* triplication and a 2-copy isogenic mutation-corrected control. *SNCA-intermediate* neurons were comparable in expression levels to human cortex samples and 2-copy *SNCA* neurons. Notably, however, normalization here is to Gapdh. We previously showed that this underestimates true neuronal αS content^58^ by several-fold in the human brain (where there are co-resident glia). *SNCA-high* neurons have approximately 2-fold higher protein levels than 4-copy *SNCA* neurons, thus likely a closer surrogate for true neuronal expression levels in human brain (Figure 3B). qPCR in DIV28 neurons confirmed *SNCA* expression at the mRNA level in the CiS models (Figure 3C).

To investigate cytotoxicity phenotype in CiS neurons, we performed longitudinal imaging (Incucyte) with a cytotox assay. Significant cell death was observed in *SNCA-high* neurons starting from DIV14 (Figure 3D). At DIV28, significantly increased Caspase+ cells were observed in *SNCA* overexpression neurons (Supplementary Figure 3A). We next investigated whether altered αS levels and resultant toxicity at DIV28 in the CiS models was accompanied by altered expression of the *PSEN2*/Aβ-actin gene module described in Figure 2. qPCR expression-analysis of key members of this module in our CiS models at DIV28 revealed that altering αS levels did significantly alter expression of these genes (Figure 3E). These data suggested that alteration of the actin cytoskeleton may be accompanying toxicity in these models.

We previously showed that αS closely interacts with Spectrin^36,37^ and can induce profound and consequential alterations in the actin cytoskeleton in *Drosophila*^62^. This earlier work demonstrated altered stabilization of actin into F-actin. Phalloidin is a dye that can recognize actin stabilization. We thus performed phalloidin immunostaining in CiS neurons. We observed no phalloidin intensity difference among DIV14 CiS neurons, but by DIV28 we observed that phalloidin intensity trended significantly lower in *SNCA-high* neurons and phalloidin aggregates were noted to form (Figure 3F-G). Thus, stabilized actin is reduced overall as αS becomes toxic.

Since our significant top-hit genes were either positive regulators or positive effectors of RhoA signaling (Figure 2A), we investigated the consequences of *RhoA* reduction in neurons as a proxy for the damaging/missense mutations identified in our study. We previously established a lenti-shRNA construct targeting *RhoA* and confirmed knockdown in both HEK cells and neurons^58^. We knocked down *RhoA* in DIV0 CiS neurons. By DIV 14, *RhoA* downregulation significantly decreased phalloidin in all CiS neurons (Figure 3H, Supplementary Figure 3B), reminiscent of the *SNCA* phenotype in DIV28 *SNCA*-*high* neurons. A standard marker of pathologic αS is phosphorylation at Ser129 (pS129), the accumulation of which occurs across all synucleinopathies. Even at day 14, where there is no overt toxicity in the DIV28 *SNCA-high* neurons, these neurons had elevated levels of this marker compared to *SNCA-intermediate* or *SNCA-endogenous* lines. After *RhoA* downregulation, αS-pS129 selectively increased in the *SNCA-high* neurons but not *SNCA-intermediate* or *SNCA-endogenous* lines (Figure 3I, Supplementary Figure 3C). Thus, in cells primed for αS-induced cell death, *RhoA* downregulation further exacerbates synucleinopathy.

### Diffused synucleinopathy in *PSEN2*/*ARHGEF1*/*ARHGEF28* mutation carriers regardless of the extent of concomitant AD pathology

The advantage of our cohort is that a subset had matched postmortem brain and highly characterized neuropathology. Since RhoA inhibition induced pathologic αS induction, we next probed more deeply into the localization of this type of pathology in carriers of mutations in *PSEN2*/Aβ-actin genes for whom we had postmortem brain samples. Among these six brains, we noted that all but one had diffuse brainstem and cortical synucleinopathy. One (*ARHGEF1* mutation carrier) had brainstem-predominant disease but we note he was an outlier among these cases with early age of onset (46yo) (Figure 4A, Figure 1D and Supplementary Figure 1). For example, the clinical phenomenology may have been attributable to some secondary pathology, especially concomitant AD. This was particularly pertinent to investigate for *PSEN2* mutation-carriers because this is a known AD gene.

Neuropathologic criteria for AD are set with three “scores” corresponding to Amyloid/Thal, Braak/tau and CERAD/neuritic plaque quantitation. Based on these scores, cases are designated as having no, low, intermediate, or high level of Alzheimer’s disease neuropathologic change (ADNC). When we examined the brains matched to our genes of interest, we found that *PSEN2*, *ARHGEF1* and *ARHGEF28* variant carriers could exhibited diffuse αS pathology completely independent of AD co-pathology (Figure 4A; Figure 1D and Supplementary Figure 1). As a case in point, we present the brain histopathology of a patient who harbored mutations in both *PSEN2* and *ARHGEF1* (Figure 4B). These findings are far from definitive, given the small numbers, but are consistent with this module of genes being important across different neuronal subtypes, including cortical and DA neurons, and contributing to diffuse brainstem and cortical synucleinopathy.

### Dysregulated expression of *PSEN2*/Aβ-actin gene module in cortical and DA neurons

Our neuropathologic data suggested there could be diffuse dysregulation of *PSEN2*/Aβ-actin gene module across different cell types in the brain. Moreover, there was enrichment of DA neuron-specific eQTLs near *PSEN2* (Supplementary Table 4) and a suggestive link with PD GWAS (Supplementary Figure 3C). This raised the possibility that even in sporadic PD/LBD cases there could be dysregulation of this gene module. To address this, we first confirmed that there are indeed cell-type specific cis-eQTLs near our *PSEN2*/Aβ-actin module member-genes in different neuronal subtypes (Figure 5A).

**Figure 5.**
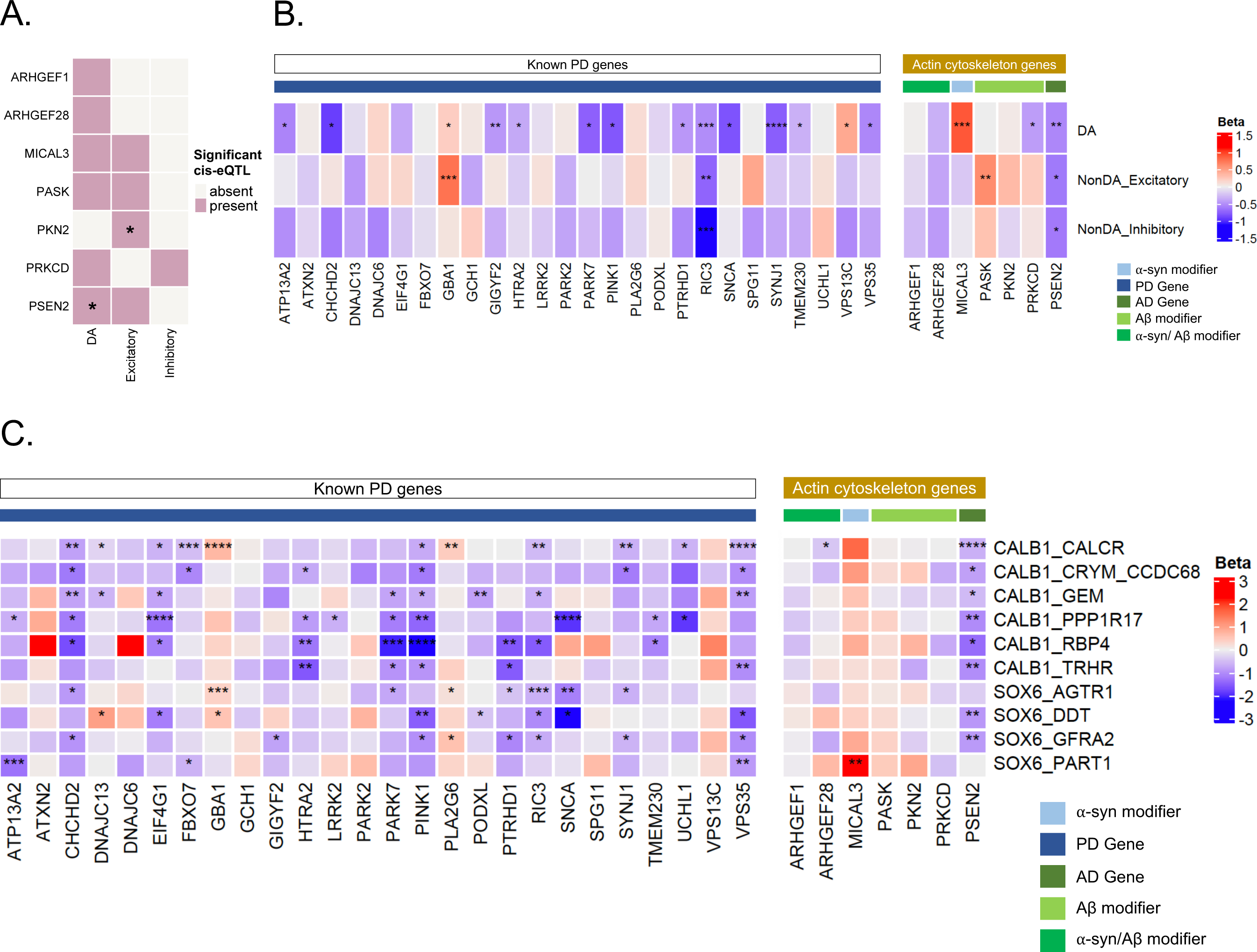
*PSEN2*/Aβ-actin gene module exhibit significant dysregulation in neuronal subpopulations in postmortem PD/DLB brain. (A) Heatmap showing the presence of cell-type-specific cis-eQTLs near the members of the *PSEN2*/Aβ-actin gene module in different neuronal cell types. *PSEN2* had a significant enrichment of DA-specific cis-eQTLs (hypergeometic p-value: 2.49×10^-65^) and *PKN2* had an enrichment of cis-eQTLs in excitatory neurons (hypergeometric p-value: 1.97×10^-77^). See Supplementary Table 4 for additional details. (B) Heatmap showing PD-associated transcriptional changes of the identified *PSEN2*/Aβ-actin gene module in post-mortem PD/DLB brains vs controls in different neuronal cell types. Cell-type specific differential expression analysis was performed in dopaminergic (DA), non-DA excitatory and non-DA inhibitory neurons of the midbrain using MAST’s discrete hurdle model. Transcriptome-wide significance is reported using a BH-corrected p-value cutoff of 0.1 (denoted by *s; *: adj-p < 0.1, **: adj-p < 0.01, ***: adj-p < 0.001, ****: adj-p < 0.0001). Transcriptional changes of 26 known PD genes were shown on the left for comparison. See Methods for details. (C) Heatmap showing PD-associated transcriptional changes of the identified *PSEN2*/Aβ-actin gene module in post-mortem PD/DLB brains vs controls in different subtypes of DA neurons in the midbrain. Cell subtype specific differential expression analysis was performed using MAST’s discrete hurdle model. Transcriptome-wide significance is reported using a BH-corrected p-value cutoff of 0.1 (denoted by *s; *: adj-p < 0.1, **: adj-p < 0.01, ***: adj-p < 0.001, ****: adj-p < 0.0001). Transcriptional changes of 26 known PD genes were shown on the left for comparison.

Next, we performed transcriptome-wide single-cell differential expression analysis using the MAST framework on the major CNS cell types using a publicly available single nuclei RNA-seq case-control dataset^63^ of ∼450k cells from ten PD and DLB patients and eight healthy controls (Methods). For comparison, we also looked at the expression changes of 26 genes that harbor either familial or common variants known to confer increased risk of PD^64^ as suggested by the original study^63^. Like known PD genes, our SKAT-O significant genes showed a diverse profile of transcriptome-wide significant (BH-corrected p-value < 0.1) expression changes across different cell types in PD brains compared to controls (Supplementary Figure 5A). Among our top hits, the known PD gene *GBA1* was significantly upregulated in dopaminergic (DA) and glutamatergic (non-DA excitatory) neurons. In contrast, the known AD gene *CR1* was downregulated considerably in microglia.

With specific reference to our PSEN2/Aβ-actin module (Figure 5B, right), *PSEN2*, *MICAL3*, and *PRKCD*, showed transcriptome-wide significant expression changes in DA neurons and two were significant in excitatory neurons in PD brains compared to controls (Fig 4A). Only one gene in this group exhibited transcriptome-wide reduction in both excitatory glutamatergic and DA neurons, and that was *PSEN2*. Most strikingly, *PSEN2* was significantly downregulated in eight out of the ten DA neuron subtypes, including CALB1_GEM and CALB1_TRHR subtypes that are relatively protected in PD (Figure 5C, right). This pattern was highly reminiscent of other PD genes (Figure 5C, left). Transcriptional changes of all SKAT-O significant genes are available in Supplementary Figure 5B.

### Genome-scale CRISPR screens reveal the essentiality of *PSEN2* in both cortical glutamatergic and DA neurons

The dysregulation of genes among our top hits and the *PSEN2*/Aβ-actin gene module in both excitatory and DA neurons (Figure 5) raised the possibility that this set of genes is important for the viability and function of these neurons. In the case of *PSEN2*, there was strong downregulation across multiple cell types (Figures 5B and 5C). We thus addressed what the consequence of this downregulation might be. Recently, the emergence of CRISPR/Cas9 technology has enabled genome-scale forward genetic screens, even in neurons and glia^65,66^. We utilized this technology to ask in an unbiased fashion: on which genes does survival of cortical and dopaminergic neurons depend?

We thus performed essentiality screens in human ESC-derived neurons. Briefly, gRNAs representing 19,993 genes were transduced into H9 hESC lines that had been knocked in for a doxycycline-inducible Cas9 at the *AAVS1* safe-harbor locus. At an early timepoint (DIV20 for cortical neurons and in the neural progenitor stage for DA neurons) Cas9 was induced, and a gRNA representation was obtained. Cells grown in parallel were aged to DIV65 (cortical neurons) or DIV42 (DA neurons) and gRNA representation again assessed to look for dropout. If gRNAs representing specific genes consistently dropped out, that gene was considered essential for the neuronal subtype (Figure 6A). Top-hit genes from our targeted exome screen were in fact enriched for dropouts in both cortical (hypergeometric test; *p*=3.9×10^-4^) and DA neurons (hypergeometric test; p=0.039). Notably, *PSEN2* proved essential for survival in both neuronal populations. Thus, its downregulation in the context of synucleinopathy (as in Figures 5B and 5C) is likely to impact viability in these subclasses of neurons.

**Figure 6.**
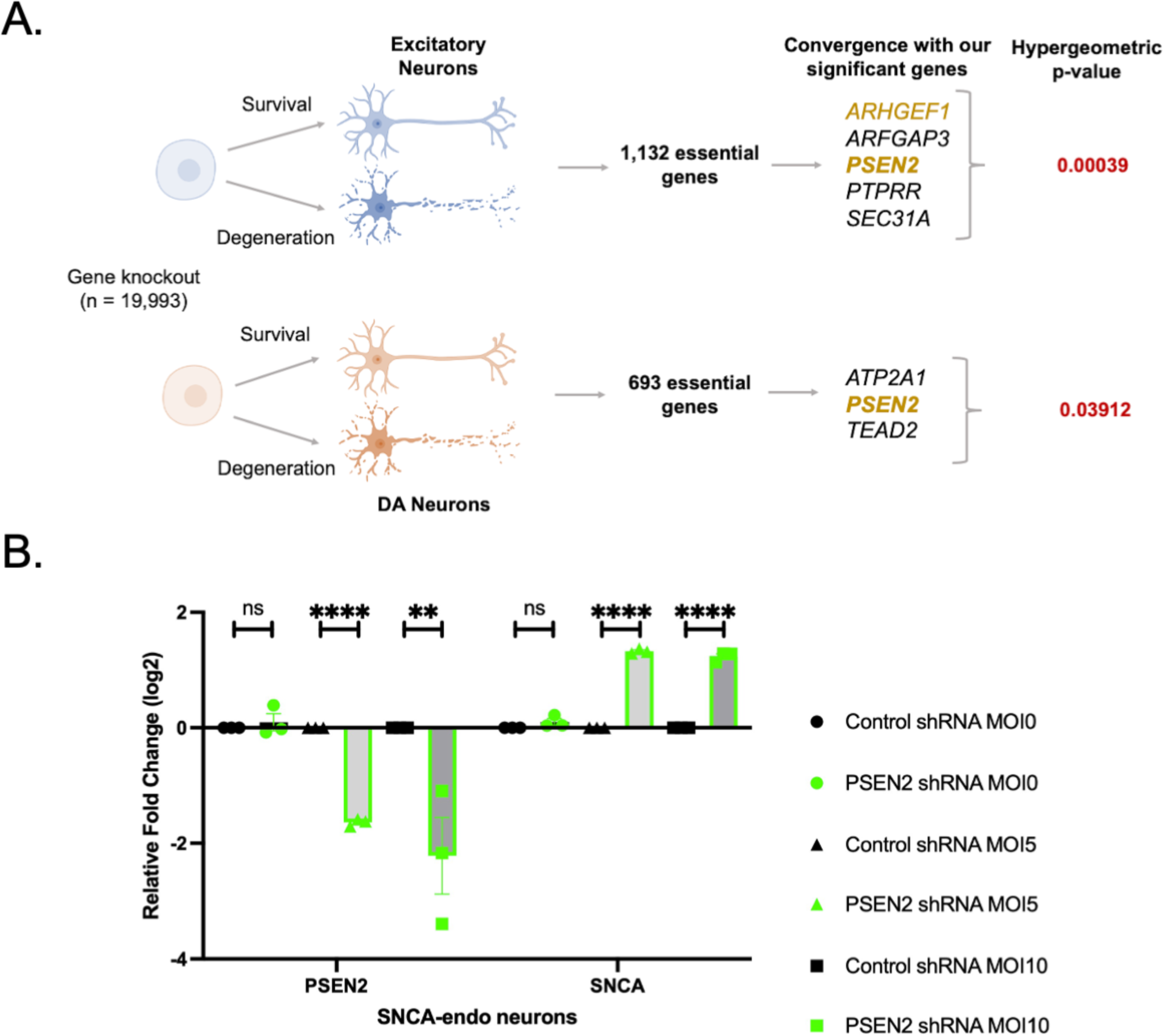
*PSEN2* is an essential gene in cortical and dopaminergic neurons, and is involved in *SNCA* expression. (A) Independent essentiality screens performed in cortical excitatory and midbrain DA neurons converge on *PSEN2*/Aβ-actin genes. In cortical and DA neurons, genome-wide screens were performed to identify genes that lead to cell death with 19,993 genes plus 1,000 control gRNAs. Screen performed in triplicate in parallel and were collected at days 0, 20, and 40 (DA neurons)/65 (cortical neurons). Samples from day 20 were compared to day 40/65 to identify barcodes with reduced representation during neuronal maturation, suggesting those genes were essential for neuronal survival. MAGeCK-MLE analysis identified 1,132 and 693 genes to be essential for cortical and DA neurons respectively. Our SKAT-O significant genes showed significant overlaps with the essential genes in hypergeometric enrichment test with *PSEN2* being the convergent signal. (B) Relative mRNA expression level in *SNCA*-endogenous CiS neurons 7days after control or *PSEN2* shRNA transduction, revealing *PSEN2* knockdown significantly increased *SNCA* expression. Values represent mean ± s.e.m. (**p < 0.01; ****p < 0.0001; t-test).

### *PSEN2* is involved in *SNCA* gene-expression

Our neuropathologic studies suggested synucleinopathy was occurring in *PSEN2* carriers independent of overt AD pathology (Figure 4). Because *PSEN2* was downregulated in synucleinopathy brains (Figures 5B and 5C), we explored whether there was any consequence of this on *SNCA* expression. We downregulated *PSEN2* in *SNCA*-endogenous neurons with shRNA. *SNCA* expression was significantly increased (Figure 6B), suggesting *PSEN2* may play an important role in regulating *SNCA* expression. Thus, one mechanism through which *PSEN2* mutation or depletion may relate to synucleinopathy is through altered regulation of *SNCA* expression. This may be particularly detrimental in cortical and DA neurons, in which *PSEN2* is both downregulated (Figures 5B and 5C) and essential for survival (Figure 6A).

## DISCUSSION

Clinical labels such as “Parkinson’s disease” or “Lewy body dementia” belie the highly heterogeneous nature of neurodegenerative diseases and the presence of abundant mixed pathologies in the brains of patients. Adopting a more molecular view of neurodegenerative disease diagnosis is likely an important first step to more targeted disease-modifying therapies for patients. In this study, rather than define patients in terms of clinical diagnoses, we anchored our genetic analysis on the presence or absence of a specific proteinopathy, in this case studying approximately 500 patients with alpha-synucleinopathies. We were motivated by recent studies suggesting that common genetic risk factors like *GBA1* exist among synucleinopathies^12,13^, from PD to DLB to MSA, and are also shared among diseases defined by different proteinaceous neuropathologies^21–23^. For example, *APOE, BIN1* or *TMEM175* may be shared risk factors among AD, PD and DLB. It thus seemed reasonable and important to conduct a genetic study directed at the presence of a proteinopathy as opposed to a single clinical diagnosis.

Human genetic investigation of neurodegeneration is limited by statistical power. For example, in recent years, adding many thousands of genomes to PD analyses has added marginal novel insights into their genetic basis. Moreover, most of the studies have investigated common genetic variation, and the recovered risk factors are predicted to have low effect. Polygenic risk analyses suggest most heritability for these diseases yet remains undefined despite decades of such investigations. Part of this missing heritability could arise from rare coding variants in the genome. But the statistical challenge for rare-variant studies is even starker, with current estimates suggesting millions of genomes will be required to adequately powered studies^67^. In this investigation, we thus focused on enhancing statistical power of our rare-variant analysis by decreasing the genomic search space on biological grounds. Simply put, the idea was to yolk unbiased forward genetic screens in model organisms with targeted human genetic screens. Saturated forward genetic screens for yeast Aβ and ɑS proteotoxicity modifiers had recovered known AD and PD risk factors at much higher rates than expected by chance. We thus hypothesized that these Aβ and ɑS cytotoxicity-modifier genes may be enriched in genetic factors of relevance to human disease. Acutely aware of the evolutionary distance between yeast and human, let alone between a unicellular organism and the human brain, we screened at high (100×) depth the exomes of patients for a wide array of homologs of each yeast hit. The idea was to survey the maximum diversity across patients, diseases, and the CNS cellular landscape.

Among the resultant significant hits at gene- and variant-level, we recovered *LRRK2* and *GBA1* as the strongest hits. This finding serves as a validating positive control because of the prior strong association between these genes and synucleinopathy risk. Second, all our hits were shared between PD and DLB diagnoses and, despite only 24 of our 496 sequenced cases having MSA, three of our top hits at the variant level (*GBA1, PTPRS* and *TEAD2*) occurred in pathologically confirmed MSA cases. Thus, our study uncovered evidence of shared risk across synucleinopathies. Moreover, disease-associated *LRRK2* and *GBA1* variants colocalized with other rare variants that reached screen-wide significance of their own accord across our cohort, raising the possibility of cumulative effects. There were sufficient *GBA1* N409S mutation carriers among both cases and controls to recover several novel genes (*PTPRB, MICAL3, FREM2, ATP2A3, SH3RF3, PRKCD*) in which variants were selectively enriched in cases, suggesting these genes may be candidates for altering phenotypic penetrance of the *GBA1* mutation. Significant hits at both variant- and gene-level originated in both Aβ and ɑS proteotoxicity networks and surprisingly included three known AD genes *CD33*, *CR1* and *PSEN2*. Even more surprisingly, top-hits from the AD/Aβ network validated in both AMP-PD and UK biobank more convincingly than for the PD/ɑS network and showed suggestive (albeit not definitive) eQTL enrichment signals among PD GWAS hits (at 1×10^-7^ significance). These data tie our genes not only to rare causal variants but to more common sporadic forms of PD also. More convincingly, our top-hit genes exhibited strong changes in expression across multiple cell types in single-cell sequencing analyses of sporadic PD/synucleinopathy brains, just as is seen with known PD genes.

Where do the Aβ and ɑS proteinopathies converge? An interesting early clue came in our prior yeast studies. While there were negligible common genetic factors among an initial genome-wide modifier screens in yeast, a subsequent pooled screen revealed a convergence at the level of *ROM1/2*, the guanidine exchange factors (GEFs) for the Rho GTPase Rho1^36^. In humans, these GEF proteins include Arhgef1 and Arhgef28. The most canonical function of these proteins is in the regulation of actin cytoskeletal dynamics, suggesting this could be a common convergent point. Indeed, actin cytoskeletal regulation emerged as a shared function of many of our top-hit genes (*ARHGEF1, ARHGEF28, PKN2, MICAL3, PRKCD*). Consistent with this, considerable emerging evidence point to abnormal stabilization of F-actin accompanying Aβ- tau- and ɑS - induced proteotoxicity and leading to altered mitochondrial dysfunction^62^. Recently, we showed a cellular aspect of synucleinopathy that could render patients particularly vulnerable to deficits in this pathway. We found that RhoA is sequestered into highly toxic classes of ɑS inclusions and that postmitotic neurons are exquisitely sensitive to RhoA reduction^58^. A similar sequestration has been described in neurofibrillary tangle pathology. One of other hits *ARGHEF28* (also known as *RGNEF*) has been implicated in ALS with sequestration into TDP-43-positive inclusions^68,69^. In this study, RhoA depletion in our novel iPSC-based “CiS” model led to F-actin destabilization in neurons and accumulation of pathologic pS129 ɑS. Collectively, these studies suggest that genetic liabilities in the RhoA signaling pathway would leave neurons particularly vulnerable to the alpha-synucleinopathy, especially those that form RhoA-positive ɑS inclusions.

The connection of *PSEN2* to synucleinopathy is particularly intriguing, given the known prominent synucleinopathy that is known to accompany *PSEN1/2*-related AD^4,70^. Importantly, the defining feature in patients in our study was synucleinopathy and not concomitant AD pathology, suggesting that the gene can be tied to a pure synucleinopathy itself. Moreover, the pathology in these patients was diffuse, affecting cortical and DA neurons. More than any other gene we recovered in this study, *PSEN2* showed the most robust transcriptional expression changes across different neuronal populations in a fashion highly reminiscent of known PD genes, including across DA subtypes (Figures 5B and 5C). This downregulation is presumably highly consequential because our genome-scale CRISPR screens pinpointed *PSEN2* as the one gene in our study that was essential for survival of both cortical glutamatergic and DA neurons. Finally, we showed that downregulation of *PSEN2* leads to a surprisingly robust elevation of *SNCA* expression in neurons. Whether or not this upregulation directly leads to neuronal toxicity (and contributes to the dropout in our CRISPR/Cas9 screens) awaits further exploration. Another open question is whether, in the specific context of cortical and DA neurons, Presenilin-2 directly regulates the actin cytoskeleton. Presenilin-2 is known to bind the filamin class of actin-binding proteins^71^. Intriguingly, ɑS itself binds filamins and many other actin-binding proteins and regulators^37,62,72^ and is in a complex with *RhoA*^58^. These are tantalizing connections for future studies to explore.

Beyond the actin cytoskeleton, altered intracellular signaling pathways were strongly represented among top hits, including a kinase within the MAPK family (*MAP2K3*), three tyrosine phosphatases strongly tied to MAPK signaling (*PTPN18*, *PTPRB*, *PTPRR*^73^, *PTPRS*) and a key transcriptional effector of Hippo signaling (*TEAD2*). Hippo signaling itself regulates MAPK signaling, and these pathways all impact cellular senescence. Importantly, *PTPRH* was tied to PD in a prior exome screen^74^ of an independent population but we did not sequence that gene here. Cellular senescence is triggered by ɑS pathology in astrocytes and microglia^75^ and by the *LRRK2*-G2019S mutation in neural cells^76^. This is an area of intense investigative focus, given the important role of aging as a risk factor across proteinopathies. Another pathway we very recently tied directly to synucleinopathy through extensive genetic and biochemical analysis is mRNA decapping and gene regulation^52^. While P-body genes did not emerge in the over-expression screen used as the source of ɑS genetic modifiers in this study (but rather in later screens), we did in fact sequence a single P-body gene, the nuclease-encoding *XRN1*, because it had emerged as a modifier of Aβ toxicity in an overexpression screen. *XRN1* was a top hit at the gene level. The emergence here not only reinforces the tie-in to synucleinopathy in our prior study, but also implicates altered mRNA stability as a potentially common shared pathway across Aβ and ɑS proteinopathies worthy of future exploration.

One key aim of our study was to investigate where in the CNS our genetic hits are impacting cellular vulnerability to neurodegeneration. Our expectation was that interactions within a unicellular context (i.e., yeast) would play out in a multitude of CNS cellular contexts. Moreover, we envisaged that these interactions could become accessible through high-depth sequencing of multiple orthologs of a single gene. In this regard, it was first reassuring that, across approximately 1500 genes in our database of Aβ- and ɑS-interacting proteins, expression of our top-hit genes was enriched in the CNS, suggesting their relevance to CNS disease. Intriguingly, when the essentiality CRISPR screen was performed in hiPSC-derived glutamatergic neurons, rather than DA neurons, a different set of essential genes emerged. It Is plausible that our top-hit genes alter vulnerability in specific cell types and may confer differential vulnerability in synucleinopathies. For example, *ARFGAP3* is a top hit in our targeted exome screen and a gene essential for survival of cortical but not DA neurons. It is tempting to speculate that mutations in a gene like *ARFGAP3* could “flip” a patient from a PD (DA>glutamatergic) to DLB (glutamatergic>DA) phenotype. More generally, our ability to uncover altered expression of genes from a study looking for rare protein-coding variants points to crossover among rare and common genetic risk factors for neurodegenerative diseases. This has already been noted for key PD-associated genes like *GBA1*, *LRRK2* and *SNCA* itself.

It will be likely decades before we have the millions of genomes required for truly systematic analysis of late-onset complex diseases. Until that time at least, and perhaps even after that time, carefully crafted biological tools will be essential to interpret and expand the reach of statistical genetics analyses. Here, our analyses suggest convergent mechanisms are shared within the synucleinopathy group, but also across proteinopathies and clinical diagnoses. This targeted rare-variant approach may effectively find true signals under the GWAS peaks for common variants. Ultimately, in this approach it is the convergence of multiple orthogonal lines of evidence that makes our gene associations convincing and are required to mitigate against false positives from biologically driven (and hence biased) studies like this. We anticipate that, among the many approaches now available, cross-species approaches will continue to yield important insights for complex diseases in the post-genomics era.

## METHODS

### Targeted exome sequencing and joint calling

We performed targeted exome sequencing of 506 familial and sporadic PD, LBD, and MSA cases as well as 98 HapMap CEU samples that were sequenced as part of the 1000 Genomes Project for quality control (Coriell Institute for Medical Research, Camden, NJ, USA). Exome enrichment and sequencing library preparation were done following a hybrid selection protocol^77^ using Agilent SureSelect baits targeting the coding region of 430 genes (∼1 Mb). Samples were sequenced at the Broad Institute of MIT and Harvard (Cambridge, MA, USA) on the Illumina HiSeq platform with ∼100× depth. Raw reads were aligned to the human reference genome (GRCh37/hg19) using BWA (v0.5.9). We used 2,570 individuals over the age of 70 who have no history of cancer and dementia from the Medical Genome Reference Bank (MGRB)^44^ as controls. Target regions were extracted from the whole genomes of these individuals with 100 bp padding around each interval, and BAM files were generated by aligning them to the same reference genome.

Individual-level variant calling was performed using the GATK HaplotypeCaller (v3.7), and GenomicsDBImport was used for joint calling genotypes of cases and controls. Variant Quality Score Recalibration (VQSR) was not performed due to the insufficient coverage of the genome by our target genes. Instead, we ensured the quality of our targeted sequencing by comparing the SNV calls from our internal HapMap CEU samples with those from the 1000 Genomes Project and observed a concordance greater than 99.9% for all chromosomes.

### Variant quality control

Variants were filtered by applying GATK’s VariantFiltration utility with the following hard filters: QD > 2.0, SOR < 3.0, MQ > 40.0, FS < 60, MQRankSum > −12.5, ReadPosRankSum > −6.0 and indels: QD > 2.0, SOR < 10.0, FS < 200.0, ReadPosRankSum > −10.0) with Q > 50 and GQ > 20. Only variants that passed all GATK filters and that overlapped our target regions were retained (Supplementary Figure 6). Variants were further filtered to include only autosomal biallelic sites with a maximum of 5% missing genotypes, HWE *p*-value > 1×10^-6^ in controls (internal HapMap and MGRB) and covered at >= 10× depth in at least 95% of samples. Sites with a depth greater than twice the mean depth were excluded, resulting in a total of 15,655 variants (15,237 SNPs and 418 indels). MGRB controls had nearly twice as many singletons (cohort allele count, AC = 1) as cases but roughly the same percentage (∼60%) as observed in gnomAD individuals with non-Finnish European (NFE).

Variants were annotated with SeattleSeq138, VEP (v95)^77^, and SnpEff (4.3t)^78^, and REVEL^79^, and rsIDs were updated to dbSNP build 150. Finally, synonymous and nonsynonymous (i.e., missense, splice, stop gained, stop lost, start lost) SNVs with a minor allele frequency (MAF) < 1% in cases, 98 internal HapMap CEU controls, gnomAD, and MGRB controls were retained.

### Sample quality control

Relatedness statistics were calculated using VCFtools (0.1.14) to determine the unadjusted A_jk_ statistic^80^. Expected value of A_jk_ is between zero and one, zero indicating unrelated individuals and one indicating duplicate samples. Based on known kinships, pairs of samples with A_jk_ > 0.3 were termed as “related.” We detected four related pairs; only one sample from each pair was retained. Additionally, we removed two cases due to excessive heterozygosity and one due to excessive genotype missingness. Principal component analysis (PCA) was performed to determine the ancestry of our cases and controls using 1397 HapMap samples from the 1000 Genomes project as reference. Three cases and 53 MGRB controls were removed as population outliers with potentially non-European ancestry. The remaining 496 cases clustered closely with the CEU (Utah residents with Northern and Western European Ancestry) and TSI (Toscani in Italia) HapMap populations and with the MGRB control samples (Supplementary Figure 7). One MGRB control was removed due to an excessive missingness rate (> 25%) which left us with 2516 MGRB controls. No additional samples were removed based on the distribution of Ts/Tv, and Het/Hom (Supplementary Figure 8).

To reduce the potential for technical artifacts, we checked whether ultra-rare variants, such as singletons (AC=1), doubletons (AC=2), and tripletons (AC=3) in cases and controls follow an exact binomial distribution (Supplementary Table 5). As singletons showed significant distributional differences between the two groups, we excluded them from our rare variant analysis.

### Rare variant association analysis

Single variant association testing was performed on rare variants (MAF < 0.01) using a one-sided Fisher exact test. The linkage between the most significant variants was assessed using LDSC, and no linkage was found.

To perform gene-level rare variant collapsing analysis, we used the CMC approach to collapse the variants. Then, we applied the SKAT-O test, a combined collapsing and variance component test, which is statistically efficient regardless of the direction and effect of the tested variants^81^. Qualifying variants were selected using an MAF cutoff of 1% and grouped using two masks: (i) nonsynonymous (missense, nonsense, and splice), and (ii) synonymous. The synonymous mask was used as an internal control.

Due to the discrete nature of rare variants and the non-uniformity of the unadjusted p-values, we performed a modified FDR adjustment^82^ for multiple hypotheses testing correction. The test performs permutation-based resampling to empirically estimate the null distribution and then uses a rank-based approach like Benjamini-Hochberg adjustment to calculate the FDR-adjusted p-value.

### Controlling for inflation

Since cases and MGRB controls were sequenced at different depths (median coverage of cases ∼55× and MGRB ∼37×), there was potential for inflation in the variant-level analysis. To check for such bias, we downsampled our cases to 1 million reads per sample (median coverage∼42X). A one-sided Fisher’s exact test of the downsampled call set detected all significant variants that were identified using the original joint call set. The genomic inflation factors in the jointly genotyped call set, the combined separately genotyped call set and the downsampled callset were computed after the variant-level association analysis. Before excluding ClinVar pathogenic mutations, the λ values were 1.32, 1.33, 1.34 for the joint callset, the combined callset, and the downsampled callset, respectively. After excluding the 15 ClinVar pathogenic mutations, the corresponding λ values decreased to 1.12, 1.15, 1.15 respectively (Supplementary Figure 9). The decrease in genomic inflation indicates that it was primarily driven by the enrichment of known pathogenic mutations rather than the difference in coverage.

### Independent validation cohorts: UK Biobank and AMP-PD

We investigated two independent Parkinson’s disease (PD) case-control cohorts from the UK Biobank (UKBB), and AMP-PD (Accelerating Medicines Partnership: Parkinson’s Disease) to validate our findings from rare variant association analysis. We used ∼500K whole exome sequencing (WES) data from UKBB. Unrelated individuals of European ancestry with age over 40 years who had a ICD10 diagnosis code, G20 (data field: 41270) were selected as PD cases. We randomly sampled 6,711 controls from 167,188 unrelated European individuals of 60 years or older with no neurological diagnosis (no ICD10 code between G01-G99) in the UKBB. For rare variant analysis, only high-quality autosomal biallelic variants passing GATK best practices filters with average quality (AQ) >= 50 were retained. Data was analyzed using the DNAnexus platform.

For the AMP-PD whole genome sequencing (WGS) dataset, we first excluded any individual belonging to the genetic registry, genetic cohort, subjects without evidence of dopamine deficit (SWEDD), prodromal categories, or the AMP-LBD cohort. We also excluded the samples from the Harvard Biomarker Study (HBS) due to the possibility of overlaps with our original case cohort. High-quality autosomal biallelic variants passing GATK best practices filters with maximum 10% missingness were retained for analysis. Unrelated European individuals were included in the analysis. Additional sample outliers were removed based on Ts/Tv, Het/Hom ratios, and per-haploid SNV counts. Outliers were defined as samples which are +/- 3 standard deviations away from the mean. After performing variant- and sample-level quality filtering, the resulting cohort included 1,598 sporadic PD cases with no known *LRRK2* and *GBA1* mutations, and 1,095 neurotypical controls.

Variants from both the cohorts were called using the GRCh38 assembly and were annotated with the gnomAD minor allele frequencies and in-silico predictions of deleteriousness of the missense variants by PolyPhen2 and SIFT from the dbNSFP (v4.3a) database using the using VEP (v109). Variants termed as synonymous, missense, splice donor, splice acceptor, splice region, stop-gained, stop-lost, start-lost, frameshift, in-frame insertion, and in-frame deletion, were included in the analysis. Three masks were used to group variants: (i) Damaging: missense variants predicted to be either ‘‘P’’ or ‘‘D’’ by PolyPhen2 or ‘‘deleterious’’ by SIFT, as well as, splice donor, splice acceptor, splice region, stop-gained, stop-lost, start-lost, frameshift, in-frame insertion, and in-frame deletion; (ii) Missense, and (iii) Synonymous.

### Rare variant trend test (RVTT)

Since single-variant and gene-level analysis of both UKBB and AMP-PD were unable to find any genome/exome-wide significant gene other than LRRK2 and GBA1 in previous studies^45,46^ as well as our in-house analysis, we utilized RVTT^51^, a gene set based approach to assess trend in rare variant burden. Due to relatively small sample sizes of PD cohorts and the burden of multiple hypotheses correction, the statistical power of variant- and gene-level burden tests remains low. RVTT increases power by looking at gene sets and reduces false positives by looking at the frequency of qualifying variants in the gene set of interest instead of presence or absence of variants. Under the null hypothesis, the test assumes that there is no linear trend in the binomial proportions of cases and controls in terms of rare variant occurrences in the gene set. The alternative hypothesis indicates the presence of a linear trend. RVTT uses the Cochran-Armitage statistic to quantify this trend. Qualifying rare variants are selected using a variable threshold approach^83^. Since the test statistic doesn’t follow a chi-square distribution, p-values are drawn from an empirical distribution estimated from 10,000 random permutations of the case-control labels.

We tested four gene sets: (i) significant genes (variant-level; FDR adj-p < 0.1), (ii) significant genes (SKAT-O; FDR adj-p < 0.1), (iii) AD genes and Aβ modifiers among significant genes, and (iv) PD genes and a-syn modifiers among significant genes (Figure 1E). RVTT was applied to both UKBB and AMP-PD. The test selected qualifying rare variants by varying the MAF cutoff up to 0.01. The validity of significant results was confirmed by demonstrating that there is an accumulation of damaging and missense variants in patients versus controls, but no statistically significant difference in proportions of synonymous variants within the same gene sets in cases vs. controls.

The RVTT p-values per variant category from both datasets were combined using the Cauchy combination test (CCT)^53^. CCT defines the test statistic as a weighted sum of transformed p-values that follows a standard Cauchy distribution. Unlike Fisher’s combined test, CCT is robust against arbitrary correlation structures among the p-values being tested and less biased towards extremely small p-values from individual studies. Therefore, it is better suited to genomic analysis where complete independence of underlying variants and genes cannot be assumed.

### Single-cell transcriptomic analysis in post-mortem brain data

We analyzed publicly available single nuclei RNA-seq data from post-mortem brains of 10 PD/DLB patients and 8 healthy controls^63^. We used the cell type and subtype definitions from the original study and followed their quality control recommendations. Differential expression analysis was performed in all major cell types of the central nervous system (CNS) as well as in the ten subtypes of dopaminergic (DA) neurons using MAST (model-based analysis of single-cell transcriptomes)^84^.

We used MAST’s discrete hurdle model which is a mixed effects model that assumes the snRNA-seq data follows a mixture of a binomial and normal distribution while accounting for pre-defined covariates. Sex, age, race, percentage of reads aligned to mitochondrial genes, and number of UMIs (log-scale) were used as fixed-effect covariates. Disease status was also included as fixed-effect covariate and was the main variable of interest. Sample ids were used as a random-effect covariate, to account for dependencies between cells originating from the same individual. The beta value from Wald test was used as an estimate of the effect of disease on expression of each gene where the sign of beta indicated the direction of effect. Differentially expressed genes were selected as being transcriptome-wide significant with a cutoff of BH-adjusted p-value < 0.1.

### Enrichment analysis of expression quantitative trait loci (eQTLs)

We searched the bulk data from brain GTEx (v8) and PsychENCODE for the presence of known expression QTLs (eQTL) and splice QTLs (sQTLs) within +/- 1 Mb of our significant genes (full set as well as the AD genes and Aβ modifiers subset) in different brain regions. We assessed the enrichment of these gene sets with significant rare variant burden over all genes having e/sQTLs in both brain GTEx and PsychENCODE and within anchored Hi-C loops. Confidence intervals were empirically estimated through random sampling of N (N=number of genes of interest), permuting 1,000 times and permutation p-values were reported.

We interrogated two publicly available datasets^56,57^ through scQTLbase^85^ to identify significant cell-type specific cis-eQTLs in near our significant genes in major CNS cell types. To assess the relationship between expression and PD risk, we integrated cell-type specific eQTLs from iPSC-derived cell types differentiating towards a mid-brain fate^57^ with summary statistics from a large-scale genome-wide association study (GWAS) of PD^43^. Since current single cell eQTL datasets for CNS cell-types have small sample size, many eQTLs remain undiscovered^86^. Also, at the genome-wide significance level (p-value < 1×10^-8^), we are underpowered to detect any enrichment. Therefore, we used different relaxed p-value cutoffs (1×10^-7^, 1×10^-6^, and 1×10^-5^) for selecting PD-associated variants with suggestive evidence. We then performed enrichment analyses using cis-eQTLs in iPSC-derived neurons near our significant genes as well as the AD genes and Aβ modifiers subset over a background of cell-type specific cis-eQTLs with suggestive signal in PD GWAS. Confidence intervals were empirically estimated through random sampling of N (N=number of genes of interest), permuting 1,000 times and permutation p-values were reported.

### Human iPSCs culture and induced neuron differentiation

Human iPSCs were cultured in Essential 8 Medium (Gibco/Thermo Fisher Scientific; Cat. No. A1517001) on 6-well plates coated with Matrigel Matrix (Corning; Cat. No. 356231) diluted 1:100 in Knockout DMEM (Gibco/Thermo Fisher Scientific; Cat. No. 10829-018). Briefly, Essential 8 Medium was replaced every day. When 80% confluent, cells were passaged with StemPro Accutase Cell Dissociation Reagent (Gibco/Thermo Fisher Scientific; Cat. No. A11105-01). Human iPSCs engineered to express NGN2 under a doxycycline-inducible system in the AAVS1 safe harbor locus were differentiated following previously published protocol^65^. Briefly, iPSCs were released as above, centrifuged, and resuspended in N2 Pre-Differentiation Medium containing the following: Knockout DMEM/F12 (Gibco/Thermo Fisher Scientific; Cat. No. 12660-012) as the base, 1X MEM Non-Essential Amino Acids (Gibco/Thermo Fisher Scientific; Cat. No. 11140-050), 1X N2 Supplement (Gibco/Thermo Fisher Scientific; Cat. No. 17502-048), 10ng/mL NT-3 (PeproTech; Cat. No. 450-03), 10ng/mL BDNF (PeproTech; Cat. No. 450-02), 1 μg/mL Mouse Laminin (Thermo Fisher Scientific; Cat. No. 23017-015), 10nM ROCK inhibitor, and 2μg/mL doxycycline hydrochloride (Sigma-Aldrich; Cat. No. D3447-500MG) to induce expression of mNGN2. iPSCs were counted and plated on Matrigel-coated plates in N2 Pre-Differentiation Medium for three days. After three days, hereafter Day 0, pre-differentiated cells were released and centrifuged as above, and pelleted cells were resuspended in Classic Neuronal Medium containing the following: half DMEM/F12 (Gibco/Thermo Fisher Scientific; Cat. No. 11320-033) and half Neurobasal-A (Gibco/Thermo Fisher Scientific; Cat. No. 10888-022) as the base, 1X MEM Non-Essential Amino Acids, 0.5X GlutaMAX Supplement (Gibco/Thermo Fisher Scientific; Cat. No. 35050-061), 0.5XN2 Supplment, 0.5XB27 Supplement (Gibco/Thermo Fisher Scientific; Cat. No. 17504-044), 10ng/mL NT-3, 10ng/mL BDNF, 1μg/mL Mouse Laminin, and 2μg/mL doxycycline hydrochloride. Pre-differentiated cells were subsequently counted and plated on BioCoat Poly-D-Lysine coated plates (Corning; Cat. No. 356470) in Classic Neuronal Medium. On Day 7and each week after, medium change was performed without doxycycline was added. For the longitudinal imaging assay, cytotox green (Sartorius; Cat. No. 4633) was added in Classic Neuronal Medium at a concentration of 250nM.

### Cortical induced synucleinopathy (CiS) model Generation

SNCA-IRES-mCherry or IRES-mCherry was cloned into lentiviral vector under the control of EF-1a promoter. Plasmids were sequenced verified and submitted for lentivirus production. WTC11 hiPSCs with doxycycline-inducible NGN2 in AAVS1 safe-harbor locus^65^ were transduced either lentivirus. hiPSCs were flow cytometry-sorted utilizing the mCherry fluorescence and replated at low density for single-cell clonal selection. iPSC clones were expanded, karyotyped.

### Whole-cell protein extraction

For cell lysis, frozen cell pellets were resuspended in RIPA buffer (Thermo Scientific; Cat. No. 89900) supplemented with Complete EDTA-free protease inhibitor cocktail (Sigma-Aldrich; Cat. No. 11873580001) and PhosSTOP phosphatase inhibitor cocktail (Sigma-Aldrich; Cat. No. 4906845001). The cell pellets were kept on ice for 30 minutes and vortexed every 10 minutes to enable complete suspension of pellets to the RIPA buffer. After incubation, the cells were centrifuged at 21000g at tabletop centrifuge for 25 minutes at 4°C. The supernatant was then collected and transferred to a new Eppendorf tube. Protein concentration was measured with Pierce™ BCA Protein Assay Kit according to manufacturer’s guidelines (Thermo Fisher; Cat. No. 23225). Protein samples were mixed with 4X NuPAGE™ LDS Sample Buffer (Thermo Fisher; Cat. No. NP0007) supplemented with 40 mM TCEP Bond Breaker (Thermo Fisher; Cat. No. 77720) and boiled for 10 minutes at 65°C.

### Western blotting

For SDS-PAGE electrophoresis, NuPAGE™ 4-12% Bis-Tris protein gels were used (Thermo Fisher; Cat. No. NP0321BOX) with MOPS running buffer (Thermo Fisher; Cat. No. NP0001). The gels were transferred to PVDF membranes (Invitrogen; Cat. No. IB24002) with iBlot™ 2 Gel Transfer Device with the preset P0 setting. After the transfer, the membranes were fixed with 0.4% Paraformaldehyde in PBS for 15 minutes at room temperature with slight rocking (This step is especially critical for detection of α-Synuclein protein). The membranes were washed three times with PBS-T (PBS with 0.1% Tween-20) and blocked with blocking buffer (5% BSA in PBS-T) for 30 minutes at room temperature. Primary antibodies are diluted in blocking buffer and incubated overnight in cold room. After three times PBS-T washing, the secondary HRP conjugated antibodies are incubated with modified blocking buffer (1% BSA in PBS-T) for 45 minutes at room temperature with slight rocking. All the exposures were recorded digitally by iBright™ CL1000 Imaging System.

### RNA isolation, qPCR analysis

Total cellular RNA was isolated with Purelink RNA mini kit (Fisher Scientific; Cat. No.12183018A) according to manufacturer’s instructions. RNA was used to synthesized cDNA with the Superscript IV Vilo master mix with ezDNase enzyme (Fisher Scientific; Cat. No. 11766050). Quantitative SYBR green PCR assay was performed using Powerup SYBR green master mix (Fisher Scientific; Cat. No. A25777) following previously published protocol^87^. The fold change in gene expression was determined by the ΔΔCt method. *GAPDH* was used as a housekeeping gene.

### Immunofluorescence and microscopy

For immunofluorescent studies, cells were fixed in 4% PFA, washed with PBS + 0.1% Triton, blocked with 5% BSA in PBS + 0.1% Triton, and incubated in primary antibodies overnight at 4 °C. The following primary antibody concentrations were used: cleaved caspase, Cell Signaling Cat. No. 9661 (1:500); TUJ1, Biolegend Cat. No. 801213 (1:500); Phalloidin, Cytoskeleton Cat. No. PHDG1-A; pS129, Abcam Cat. No. ab184674 (1:1000). Appropriate fluorescent secondary antibodies (Alexa Fluor, Molecular Probes) were used at 1:1,000. Zeiss LSM-800 confocal microscope with 40X objective lens was used in capturing the fluorescence intensity of caspase, phalloidin and pS129 staining. The same confocal settings were used in scanning all the genotypes and treatment conditions. The confocal setting was optimized to avoid any crosstalk between the fluorophores. Image J was used in analyzing fluorescence intensity and cell/aggregates count. For phalloidin, 3 images were acquired, and two experimental replicates were used. For caspase-positive cell count, the whole well was analyzed. For pS129, fluorescence intensity in the cell body was measured and normalized to the intensity of DAPI. For each condition, 14 - 18 neurons were quantified from two wells.

### Genome-wide CRISPR/Cas9 screen in midbrain cortical excitatory neurons

The WGS was performed in WA-09 (H9) embryonic stem cells with doxycycline inducible Cas9 (iCas) knocked into the AAVS1 self-harbor locus as previously described (PMID: 24931489). To perform the screen H9-iCas pluripotent stem cells were infected with the Brunello whole genome human CRISPR knockout library^88^ at an MOI of 0.3-0.5 and at 1000× representation. Successfully transduced cells were selected for by adding 0.4ug/ml puromycin to the E8 medium. PSCs were differentiated to cortical neurons as previously described^89^. A DIV20 a no Doxycycline sample was harvested as the representation control for the screen. Cas9 was induced by the addition of Doxycycline (2 μg/ml) to the culture medium from DIV20-22. Neurons were cultured until DIV65 then harvested for library preparation and sequencing. Analysis was performed using the MAGECK-MLE pipeline as previously described^90^. Genes were considered essential if they had a negative beta score with a P-value of <0.05 and FDR of 10%.

### Genome-wide CRISPR/Cas9 screen in midbrain Dopaminergic neurons

The WGS was performed in WA-09 (H9) embryonic stem cells with doxycycline inducible Cas9 (iCas) knocked into the AAVS1 self-harbor locus as previously described^91^. Stem cells were transduced with the Gattinara human CRISPR pooled knockout library^92^ at an MOI of 0.3-0.5 and 1000X representation. Transduced stem cells were selected by puromycin and differentiated toward dopamine neurons until reaching the neural progenitor stage as described^93^. Cas9 expression was induced by doxycycline addition, cells were collected for our initial timepoint, and the remaining cells were differentiated. All remaining cells were collected for our final timepoint as neuron cell death began. Sample were processed for library preparation and sequenced. Sequencing reads were aligned to the screened library and analyzed using MAGeCK-MLE^90^, and hits were classified as having Wald-FDR < 0.05 and Beta < −0.58.

### Data and code availability

UK Biobank 500K WES data was accessed through application 41250 and is available through https://ams.ukbiobank.ac.uk. Whole genome data from the Accelerating Medicines Partnership Parkinson’s disease (AMP PD) is available on the AMP-PD Knowledge Platform (https://www.amp-pd.org). AMP PD – a public-private partnership – is managed by the FNIH and funded by Celgene, GSK, the Michael J. Fox Foundation for Parkinson’s Research, the National Institute of Neurological Disorders and Stroke, Pfizer, and Verily. AMP-PD investigators have not participated in reviewing the data analysis or content of the manuscript. Single nuclei RNA-seq data from Kamath et. al.^63^ is available from the Broad Institute Single Cell Portal: https://singlecell.broadinstitute.org/single_cell/study/SCP1768/ and GEO (accession no. <u>GSE178265</u>). Single cell eQTL data from Jerber et. al.^57^ can be downloaded from Zenodo: https://zenodo.org/record/4333872. All e/sQTL data from GTEx v8 can be downloaded from https://www.gtexportal.org/home/downloads/adult-gtex. All eQTL and Hi-C data from PsychEncode can be downloaded from http://resource.psychencode.org/. The source code for rare variant trend test (RVTT) is available on GitHub (https://github.com/snz20/RVTT) and Zenodo (DOI: 10.5281/zenodo.10627549). Single-variant Fisher exact test, gene-level SKAT-O, MAST analysis of snRNA-seq data, and MAGeCK-MLE analysis were performed using R packages stats, SKAT, MAST, and MAGeCKFlute respectively.

## Acknowledgement

Deep targeted exome sequencing of the original cohort was supported through Fidelity funds (FBRI: https://fprimecapital.com/fbri). The authors gratefully acknowledge funding by the Aligning Science Across Parkinson’s Initiative (ASAP) award ASAP-000472, Brigham Research Institute Director’s Transformative Award, Michael J. Fox. Foundation award 023460 and the National Institute of Health (NIH) grant R01NS109209. SN gratefully acknowledges support from the NIH grant R35-GM127131 (PI: Sunyaev) and the Australian Parkinson’s Mission. XW gratefully acknowledges support from the NIH grant T32AG000222 (PI: Yankner). We thank Dr. Christopher Cassa and Prof. Peter Kharchenko for sharing resources for the UKBB analysis and the snRNA-seq analysis in post-mortem brains respectively.

## REFERENCES

1 Goedert, M., Spillantini, M. G., Del Tredici, K. & Braak, H. 100 years of Lewy pathology. Nat Rev Neurol 9, 13–24 (2013). 10.1038/nrneurol.2012.242

2 Robinson, J. L. et al. Neurodegenerative disease concomitant proteinopathies are prevalent, age-related and APOE4-associated. Brain 141, 2181–2193 (2018). 10.1093/brain/awy146

3 Ye, H., Robak, L. A., Yu, M., Cykowski, M. & Shulman, J. M. Genetics and Pathogenesis of Parkinson’s Syndrome. Annu Rev Pathol 18, 95–121 (2023). 10.1146/annurev-pathmechdis-031521-034145

4 Leverenz, J. B. et al. Lewy body pathology in familial Alzheimer disease: evidence for disease- and mutation-specific pathologic phenotype. Arch Neurol 63, 370–376 (2006). 10.1001/archneur.63.3.370

5 Arai, Y., Yamazaki, M., Mori, O., Muramatsu, H., Asano, G. & Katayama, Y. Alpha-synuclein-positive structures in cases with sporadic Alzheimer’s disease: morphology and its relationship to tau aggregation. Brain Res 888, 287–296 (2001). 10.1016/s0006-8993(00)03082-1

6 Dugger, B. N. et al. Concomitant pathologies among a spectrum of parkinsonian disorders. Parkinsonism Relat Disord 20, 525–529 (2014). 10.1016/j.parkreldis.2014.02.012

7 Hepp, D. H. et al. Distribution and Load of Amyloid-beta Pathology in Parkinson Disease and Dementia with Lewy Bodies. J Neuropathol Exp Neurol 75, 936–945 (2016). 10.1093/jnen/nlw070

8 Chu, Y., Hirst, W. D., Federoff, H. J., Harms, A. S., Stoessl, A. J. & Kordower, J. H. Nigrostriatal tau pathology in parkinsonism and Parkinson’s disease. Brain 147, 444–457 (2024). 10.1093/brain/awad388

9 Zarranz, J. J. et al. The new mutation, E46K, of alpha-synuclein causes Parkinson and Lewy body dementia. Ann Neurol 55, 164–173 (2004). 10.1002/ana.10795

10 Whittaker, H. T., Qui, Y., Bettencourt, C. & Houlden, H. Multiple system atrophy: genetic risks and alpha-synuclein mutations. F1000Res 6, 2072 (2017). 10.12688/f1000research.12193.1

11 Fujishiro, H. et al. Diversity of pathological features other than Lewy bodies in familial Parkinson’s disease due to SNCA mutations. Am J Neurodegener Dis 2, 266–275 (2013).

12 Nalls, M. A. et al. A multicenter study of glucocerebrosidase mutations in dementia with Lewy bodies. JAMA Neurol 70, 727–735 (2013). 10.1001/jamaneurol.2013.1925

13 Rongve, A. et al. GBA and APOE epsilon4 associate with sporadic dementia with Lewy bodies in European genome wide association study. Sci Rep 9, 7013 (2019). 10.1038/s41598-019-43458-2

14 Walsh, D. M. & Selkoe, D. J. A critical appraisal of the pathogenic protein spread hypothesis of neurodegeneration. Nat Rev Neurosci 17, 251–260 (2016). 10.1038/nrn.2016.13

15 Peng, C. et al. Cellular milieu imparts distinct pathological alpha-synuclein strains in alpha-synucleinopathies. Nature 557, 558–563 (2018). 10.1038/s41586-018-0104-4

16 Shahnawaz, M. et al. Discriminating alpha-synuclein strains in Parkinson’s disease and multiple system atrophy. Nature 578, 273–277 (2020). 10.1038/s41586-020-1984-7

17 Lau, A. et al. alpha-Synuclein strains target distinct brain regions and cell types. Nat Neurosci 23, 21–31 (2020). 10.1038/s41593-019-0541-x

18 Melki, R. Role of Different Alpha-Synuclein Strains in Synucleinopathies, Similarities with other Neurodegenerative Diseases. J Parkinsons Dis 5, 217–227 (2015). 10.3233/JPD-150543

19 Jarosz, D. F. & Khurana, V. Specification of Physiologic and Disease States by Distinct Proteins and Protein Conformations. Cell 171, 1001–1014 (2017). 10.1016/j.cell.2017.10.047

20 Lam, I., Hallacli, E. & Khurana, V. Proteome-Scale Mapping of Perturbed Proteostasis in Living Cells. Cold Spring Harb Perspect Biol 12 (2020). 10.1101/cshperspect.a034124

21 Chia, R. et al. Genome sequencing analysis identifies new loci associated with Lewy body dementia and provides insights into its genetic architecture. Nat Genet 53, 294–303 (2021). 10.1038/s41588-021-00785-3

22 Reynolds, R. H. et al. Local genetic correlations exist among neurodegenerative and neuropsychiatric diseases. NPJ Parkinsons Dis 9, 70 (2023). 10.1038/s41531-023-00504-1

23 Brainstorm, C. et al. Analysis of shared heritability in common disorders of the brain. Science 360 (2018). 10.1126/science.aap8757

24 Hutton, M. Molecular genetics of chromosome 17 tauopathies. Ann N Y Acad Sci 920, 63–73 (2000). 10.1111/j.1749-6632.2000.tb06906.x

25 Strickland, S. L. et al. MAPT haplotype-stratified GWAS reveals differential association for AD risk variants. Alzheimers Dement 16, 983–1002 (2020). 10.1002/alz.12099

26 Charlesworth, G. et al. Tau acts as an independent genetic risk factor in pathologically proven PD. Neurobiol Aging 33, 838 e837–811 (2012). 10.1016/j.neurobiolaging.2011.11.001

27 Dickson, D. W. et al. APOE epsilon4 is associated with severity of Lewy body pathology independent of Alzheimer pathology. Neurology 91, e1182–e1195 (2018). 10.1212/WNL.0000000000006212

28 Henderson, M. X., Sengupta, M., Trojanowski, J. Q. & Lee, V. M. Y. Alzheimer’s disease tau is a prominent pathology in LRRK2 Parkinson’s disease. Acta Neuropathol Commun 7, 183 (2019). 10.1186/s40478-019-0836-x

29 Clinton, L. K., Blurton-Jones, M., Myczek, K., Trojanowski, J. Q. & LaFerla, F. M. Synergistic Interactions between Abeta, tau, and alpha-synuclein: acceleration of neuropathology and cognitive decline. J Neurosci 30, 7281–7289 (2010). 10.1523/JNEUROSCI.0490-10.2010

30 Kayed, R., Dettmer, U. & Lesne, S. E. Soluble endogenous oligomeric alpha-synuclein species in neurodegenerative diseases: Expression, spreading, and cross-talk. J Parkinsons Dis 10, 791–818 (2020). 10.3233/JPD-201965

31 Chia, S. et al. Monomeric and fibrillar alpha-synuclein exert opposite effects on the catalytic cycle that promotes the proliferation of Abeta42 aggregates. Proc Natl Acad Sci U S A 114, 8005–8010 (2017). 10.1073/pnas.1700239114

32 Guo, J. L. et al. Distinct alpha-synuclein strains differentially promote tau inclusions in neurons. Cell 154, 103–117 (2013). 10.1016/j.cell.2013.05.057

33 Khurana, V. & Lindquist, S. Modelling neurodegeneration in Saccharomyces cerevisiae: why cook with baker’s yeast? Nat Rev Neurosci 11, 436–449 (2010). 10.1038/nrn2809

34 Treusch, S. et al. Functional links between Abeta toxicity, endocytic trafficking, and Alzheimer’s disease risk factors in yeast. Science 334, 1241–1245 (2011). 10.1126/science.1213210

35 Yeger-Lotem, E. et al. Bridging high-throughput genetic and transcriptional data reveals cellular responses to alpha-synuclein toxicity. Nat Genet 41, 316–323 (2009). 10.1038/ng.337

36 Khurana, V. et al. Genome-Scale Networks Link Neurodegenerative Disease Genes to alpha-Synuclein through Specific Molecular Pathways. Cell Syst 4, 157–170 e114 (2017). 10.1016/j.cels.2016.12.011

37 Chung, C. Y. et al. In Situ Peroxidase Labeling and Mass-Spectrometry Connects Alpha-Synuclein Directly to Endocytic Trafficking and mRNA Metabolism in Neurons. Cell Syst 4, 242–250 e244 (2017). 10.1016/j.cels.2017.01.002

38 Ozaki, K. et al. Rom1p and Rom2p are GDP/GTP exchange proteins (GEPs) for the Rho1p small GTP binding protein in Saccharomyces cerevisiae. EMBO J 15, 2196–2207 (1996).

39 Zhang, J. et al. The association of GNB5 with Alzheimer disease revealed by genomic analysis restricted to variants impacting gene function. Am J Hum Genet (2024). 10.1016/j.ajhg.2024.01.005

40 Bandres-Ciga, S. et al. Large-scale pathway specific polygenic risk and transcriptomic community network analysis identifies novel functional pathways in Parkinson disease. Acta Neuropathol 140, 341–358 (2020). 10.1007/s00401-020-02181-3

41 Bandres-Ciga, S. et al. The endocytic membrane trafficking pathway plays a major role in the risk of Parkinson’s disease. Mov Disord 34, 460–468 (2019). 10.1002/mds.27614

42 Robak, L. A. et al. Excessive burden of lysosomal storage disorder gene variants in Parkinson’s disease. Brain 140, 3191–3203 (2017). 10.1093/brain/awx285

43 Nalls, M. A. et al. Identification of novel risk loci, causal insights, and heritable risk for Parkinson’s disease: a meta-analysis of genome-wide association studies. Lancet Neurol 18, 1091–1102 (2019). 10.1016/S1474-4422(19)30320-5

44 Pinese, M. et al. The Medical Genome Reference Bank contains whole genome and phenotype data of 2570 healthy elderly. Nat Commun 11, 435 (2020). 10.1038/s41467-019-14079-0

45 Makarious, M. B. et al. Large-scale rare variant burden testing in Parkinson’s disease. Brain 146, 4622–4632 (2023). 10.1093/brain/awad214

46 Karczewski, K. J. et al. Systematic single-variant and gene-based association testing of thousands of phenotypes in 394,841 UK Biobank exomes. Cell Genom 2, 100168 (2022). 10.1016/j.xgen.2022.100168

47 Rana, H. Q., Balwani, M., Bier, L. & Alcalay, R. N. Age-specific Parkinson disease risk in GBA mutation carriers: information for genetic counseling. Genet Med 15, 146–149 (2013). 10.1038/gim.2012.107

48 Lee, A. J. et al. Penetrance estimate of LRRK2 p.G2019S mutation in individuals of non-Ashkenazi Jewish ancestry. Mov Disord 32, 1432–1438 (2017). 10.1002/mds.27059

49 Riboldi, G. M. & Di Fonzo, A. B. GBA, Gaucher Disease, and Parkinson’s Disease: From Genetic to Clinic to New Therapeutic Approaches. Cells 8 (2019). 10.3390/cells8040364

50 Morgenthaler, S. & Thilly, W. G. A strategy to discover genes that carry multi-allelic or mono-allelic risk for common diseases: a cohort allelic sums test (CAST). Mutat Res 615, 28–56 (2007). 10.1016/j.mrfmmm.2006.09.003

51 Bendapudi, P. K. et al. Low-frequency inherited complement receptor variants are associated with purpura fulminans. Blood (2023). 10.1182/blood.2023021231

52 Hallacli, E. et al. The Parkinson’s disease protein alpha-synuclein is a modulator of processing bodies and mRNA stability. Cell 185, 2035–2056 e2033 (2022). 10.1016/j.cell.2022.05.008

53 Liu, Y. & Xie, J. Cauchy combination test: a powerful test with analytic p-value calculation under arbitrary dependency structures. J Am Stat Assoc 115, 393–402 (2020). 10.1080/01621459.2018.1554485

54 Li, J., Kong, N., Han, B. & Sul, J. H. Rare variants regulate expression of nearby individual genes in multiple tissues. PLoS Genet 17, e1009596 (2021). 10.1371/journal.pgen.1009596

55 Momozawa, Y. & Mizukami, K. Unique roles of rare variants in the genetics of complex diseases in humans. J Hum Genet 66, 11–23 (2021). 10.1038/s10038-020-00845-2

56 Bryois, J. et al. Cell-type-specific cis-eQTLs in eight human brain cell types identify novel risk genes for psychiatric and neurological disorders. Nat Neurosci 25, 1104–1112 (2022). 10.1038/s41593-022-01128-z

57 Jerber, J. et al. Population-scale single-cell RNA-seq profiling across dopaminergic neuron differentiation. Nat Genet 53, 304–312 (2021). 10.1038/s41588-021-00801-6

58 Lam, I. et al. Rapid iPSC inclusionopathy models shed light on formation, consequence and molecular subtype of α-synuclein inclusions. bioRxiv, 2022.2011.2008.515615 (2022). 10.1101/2022.11.08.515615

59 Pantazis, C. B. et al. A reference human induced pluripotent stem cell line for large-scale collaborative studies. Cell Stem Cell 29, 1685–1702 e1622 (2022). 10.1016/j.stem.2022.11.004

60 Ramos, D. M., Skarnes, W. C., Singleton, A. B., Cookson, M. R. & Ward, M. E. Tackling neurodegenerative diseases with genomic engineering: A new stem cell initiative from the NIH. Neuron 109, 1080–1083 (2021). 10.1016/j.neuron.2021.03.022

61 Gwinn-Hardy, K. et al. Distinctive neuropathology revealed by alpha-synuclein antibodies in hereditary parkinsonism and dementia linked to chromosome 4p. Acta Neuropathol 99, 663–672 (2000). 10.1007/s004010051177

62 Ordonez, D. G., Lee, M. K. & Feany, M. B. alpha-synuclein Induces Mitochondrial Dysfunction through Spectrin and the Actin Cytoskeleton. Neuron 97, 108–124 e106 (2018). 10.1016/j.neuron.2017.11.036

63 Kamath, T. et al. Single-cell genomic profiling of human dopamine neurons identifies a population that selectively degenerates in Parkinson’s disease. Nat Neurosci 25, 588–595 (2022). 10.1038/s41593-022-01061-1

64 Bustos, B. I., Krainc, D., Lubbe, S. J. & Consortium, f. T. I. P. s. D. G. Whole-exome analysis in Parkinson’s disease reveals a high burden of ultra rare variants in early onset cases. bioRxiv, 2020.2006.2006.137299 (2020). 10.1101/2020.06.06.137299

65 Tian, R. et al. CRISPR Interference-Based Platform for Multimodal Genetic Screens in Human iPSC-Derived Neurons. Neuron 104, 239–255 e212 (2019). 10.1016/j.neuron.2019.07.014

66 Tian, R. et al. Genome-wide CRISPRi/a screens in human neurons link lysosomal failure to ferroptosis. Nat Neurosci 24, 1020–1034 (2021). 10.1038/s41593-021-00862-0

67 O’Connor, L. J. The distribution of common-variant effect sizes. Nat Genet 53, 1243–1249 (2021). 10.1038/s41588-021-00901-3

68 Abbassi, Y. et al. Axon guidance genes are regulated by TDP-43 and RGNEF through the rate of long-intron processing. bioRxiv, 2023.2012.2005.570131 (2023). 10.1101/2023.12.05.570131

69 Farhan, S. M. K., Gendron, T. F., Petrucelli, L., Hegele, R. A. & Strong, M. J. ARHGEF28 p.Lys280Metfs40Ter in an amyotrophic lateral sclerosis family with a C9orf72 expansion. Neurol Genet 3, e190 (2017). 10.1212/NXG.0000000000000190

70 Geiger, J. T. et al. Next-generation sequencing reveals substantial genetic contribution to dementia with Lewy bodies. Neurobiol Dis 94, 55–62 (2016). 10.1016/j.nbd.2016.06.004

71 Zhang, W., Han, S. W., McKeel, D. W., Goate, A. & Wu, J. Y. Interaction of presenilins with the filamin family of actin-binding proteins. J Neurosci 18, 914–922 (1998). 10.1523/JNEUROSCI.18-03-00914.1998

72 Oliveira da Silva, M. I. & Liz, M. A. Linking Alpha-Synuclein to the Actin Cytoskeleton: Consequences to Neuronal Function. Front Cell Dev Biol 8, 787 (2020). 10.3389/fcell.2020.00787

73 Hendriks, W. J., Dilaver, G., Noordman, Y. E., Kremer, B. & Fransen, J. A. PTPRR protein tyrosine phosphatase isoforms and locomotion of vesicles and mice. Cerebellum 8, 80–88 (2009). 10.1007/s12311-008-0088-y

74 Jansen, I. E. et al. Discovery and functional prioritization of Parkinson’s disease candidate genes from large-scale whole exome sequencing. Genome Biol 18, 22 (2017). 10.1186/s13059-017-1147-9

75 Verma, D. K. et al. Alpha-Synuclein Preformed Fibrils Induce Cellular Senescence in Parkinson’s Disease Models. Cells 10 (2021). 10.3390/cells10071694

76 Ho, D. H., Seol, W. & Son, I. Upregulation of the p53-p21 pathway by G2019S LRRK2 contributes to the cellular senescence and accumulation of alpha-synuclein. Cell Cycle 18, 467–475 (2019). 10.1080/15384101.2019.1577666

77 McLaren, W. et al. The Ensembl Variant Effect Predictor. Genome Biol 17, 122 (2016). 10.1186/s13059-016-0974-4

78 Cingolani, P. et al. A program for annotating and predicting the effects of single nucleotide polymorphisms, SnpEff: SNPs in the genome of Drosophila melanogaster strain w1118; iso-2; iso-3. Fly (Austin) 6, 80–92 (2012). 10.4161/fly.19695

79 Ioannidis, N. M. et al. REVEL: An Ensemble Method for Predicting the Pathogenicity of Rare Missense Variants. Am J Hum Genet 99, 877–885 (2016). 10.1016/j.ajhg.2016.08.016

80 Yang, J. et al. Common SNPs explain a large proportion of the heritability for human height. Nat Genet 42, 565–569 (2010). 10.1038/ng.608

81 Lee, S. et al. Optimal unified approach for rare-variant association testing with application to small-sample case-control whole-exome sequencing studies. Am J Hum Genet 91, 224–237 (2012). 10.1016/j.ajhg.2012.06.007

82 Dietlein, F. et al. Identification of cancer driver genes based on nucleotide context. Nat Genet 52, 208–218 (2020). 10.1038/s41588-019-0572-y

83 Price, A. L. et al. Pooled association tests for rare variants in exon-resequencing studies. Am J Hum Genet 86, 832–838 (2010). 10.1016/j.ajhg.2010.04.005

84 Finak, G. et al. MAST: a flexible statistical framework for assessing transcriptional changes and characterizing heterogeneity in single-cell RNA sequencing data. Genome Biol 16, 278 (2015). 10.1186/s13059-015-0844-5

85 Ding, R. et al. scQTLbase: an integrated human single-cell eQTL database. Nucleic Acids Res 52, D1010–D1017 (2024). 10.1093/nar/gkad781

86 Mostafavi, H., Spence, J. P., Naqvi, S. & Pritchard, J. K. Systematic differences in discovery of genetic effects on gene expression and complex traits. Nat Genet 55, 1866–1875 (2023). 10.1038/s41588-023-01529-1

87 Wang, X. et al. Structural interaction between DISC1 and ATF4 underlying transcriptional and synaptic dysregulation in an iPSC model of mental disorders. Mol Psychiatry 26, 1346–1360 (2021). 10.1038/s41380-019-0485-2

88 Doench, J. G. et al. Optimized sgRNA design to maximize activity and minimize off-target effects of CRISPR-Cas9. Nat Biotechnol 34, 184–191 (2016). 10.1038/nbt.3437

89 Ciceri, G. et al. An epigenetic barrier sets the timing of human neuronal maturation. Nature 626, 881–890 (2024). 10.1038/s41586-023-06984-8

90 Wang, B. et al. Integrative analysis of pooled CRISPR genetic screens using MAGeCKFlute. Nat Protoc 14, 756–780 (2019). 10.1038/s41596-018-0113-7

91 Gonzalez, F. et al. An iCRISPR platform for rapid, multiplexable, and inducible genome editing in human pluripotent stem cells. Cell Stem Cell 15, 215–226 (2014). 10.1016/j.stem.2014.05.018

92 DeWeirdt, P. C. et al. Genetic screens in isogenic mammalian cell lines without single cell cloning. Nat Commun 11, 752 (2020). 10.1038/s41467-020-14620-6

93 Kim, T. W. et al. Biphasic Activation of WNT Signaling Facilitates the Derivation of Midbrain Dopamine Neurons from hESCs for Translational Use. Cell Stem Cell 28, 343–355 e345 (2021). 10.1016/j.stem.2021.01.005

