## Supplementary Figures 1 to 9 for "Deep sequencing of proteotoxicity modifier genes uncovers a Presenilin-2/beta-amyloid-actin genetic risk module shared among alpha-synucleinopathies"

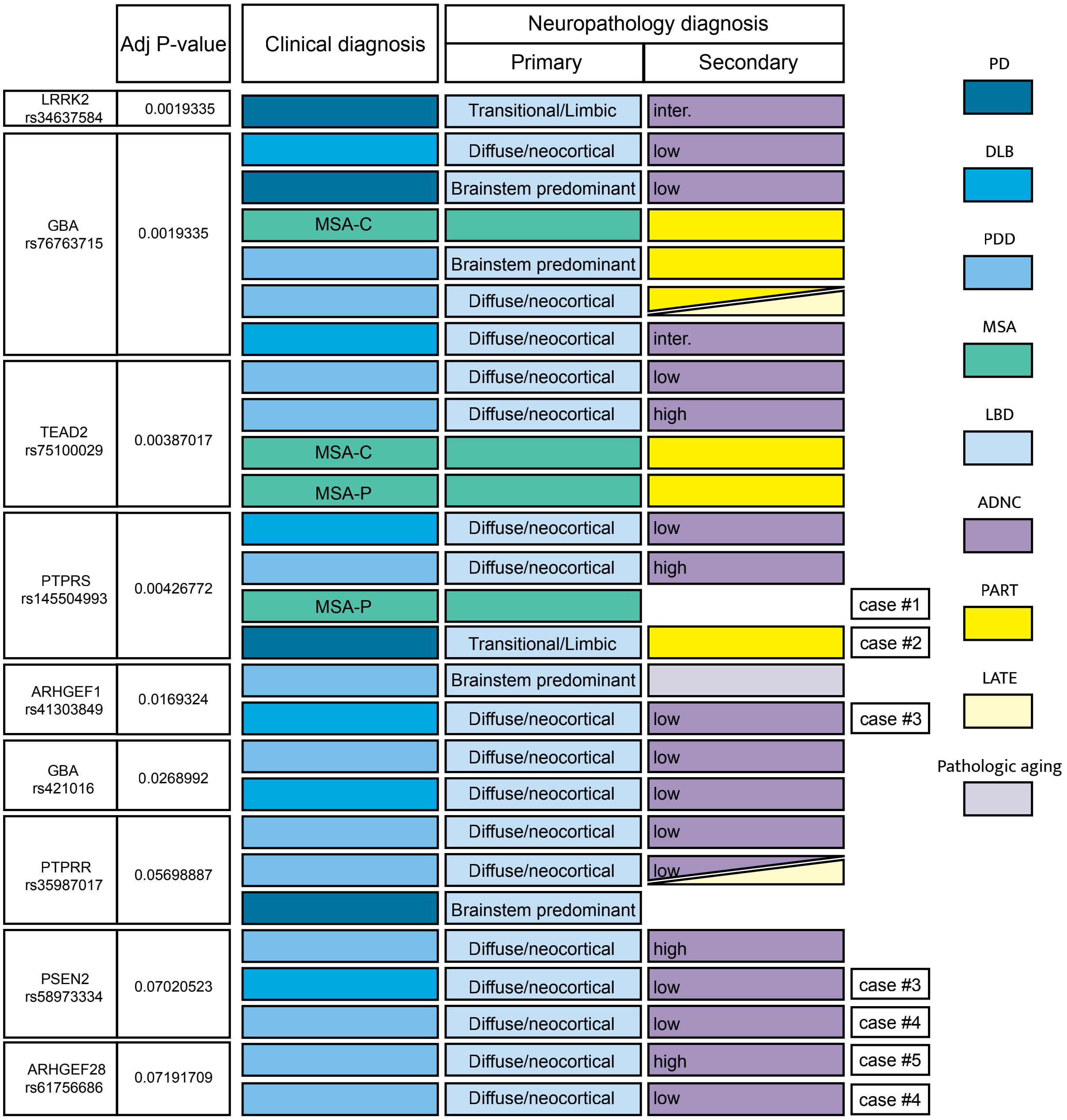

Adj P-value
Clinical diagnosis
Neuropathology diagnosis
Primary
Secondary
LRRK2 rs34637584
0.0019335
Transitional/Limbic
inter.
GBA rs76763715
0.0019335
Diffuse/neocortical
low
Brainstem predominant
low
MSA-C
Brainstem predominant
Diffuse/neocortical
Diffuse/neocortical
inter.
TEAD2 rs75100029
0.00387017
Diffuse/neocortical
low
Diffuse/neocortical
high
MSA-C
MSA-P
PTPRS rs145504993
0.00426772
Diffuse/neocortical
low
Diffuse/neocortical
high
MSA-P
case #1
Transitional/Limbic
case #2
ARHGEF1 rs41303849
0.0169324
Brainstem predominant
Diffuse/neocortical
low
case #3
GBA rs421016
0.0268992
Diffuse/neocortical
low
Diffuse/neocortical
low
PTPRR rs35987017
0.05698887
Diffuse/neocortical
low
Diffuse/neocortical
low
Brainstem predominant
PSEN2 rs58973334
0.07020523
Diffuse/neocortical
high
Diffuse/neocortical
low
case #3
Diffuse/neocortical
low
case #4
ARHGEF28 rs61756686
0.07191709
Diffuse/neocortical
high
case #5
Diffuse/neocortical
low
case #4
PD
DLB
PDD
MSA
LBD
ADNC
PART
LATE
Pathologic aging


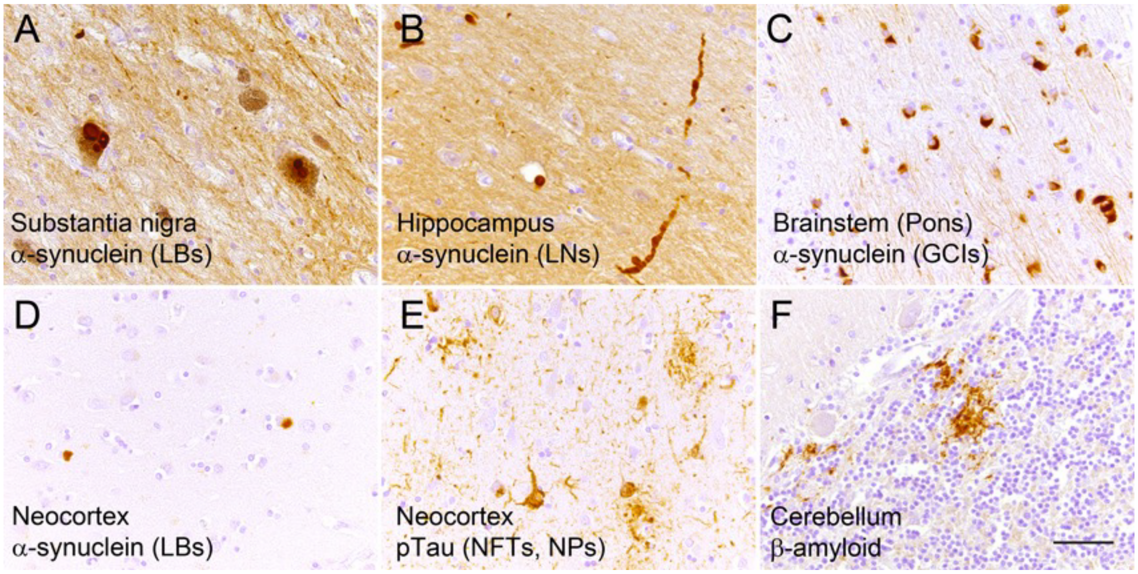

A
Substantia nigra
α-synuclein (LBs)
B
Hippocampus
α-synuclein (LNs)
C
Brainstem (Pons)
α-synuclein (GCIs)
D
Neocortex
α-synuclein (LBs)
E
Neocortex
pTau (NFTs, NPs)
F
Cerebellum
β-amyloid

**Supplementary Figure 1. Clinical and neuropathology diagnosis of patients carrying genetic variants that emerged as top hits in the association test** (cases vs MGRB control; FDR adjusted p-value <0.1). (Upper panel) ADNC: Alzheimer disease neuropathologic change; LATE: limbic predominant age-related TDP-43 encephalopathy; MCI: mild cognitive impairment; MSA-C: multiple system atrophy cerebellar type; MSA-P: multiple system atrophy parkinsonian type; PART: primary age-related tauopathy (tau deposits found without beta-amyloid); PDD: Parkinson's disease with dementia. "Pathologic aging" refers to beta-amyloid deposits being found without any tau. (A, B) case #2: relatively pure LBD (transitional/limbic pattern) in patient with PD. (C) case #1: a classic example of MSA. (D- F) case #5: LBD (diffuse/neocortical pattern) with a high level of AD neuropathologic change. (D) neocortical Lewy bodies; (E) phospho- tau stain showing neocortical neurofibrillary tangles and neuritic plaques; (F) β-amyloid deposits in the cerebellum (which only happens in cases with advance amyloid deposition) (scalebar: 50 um).

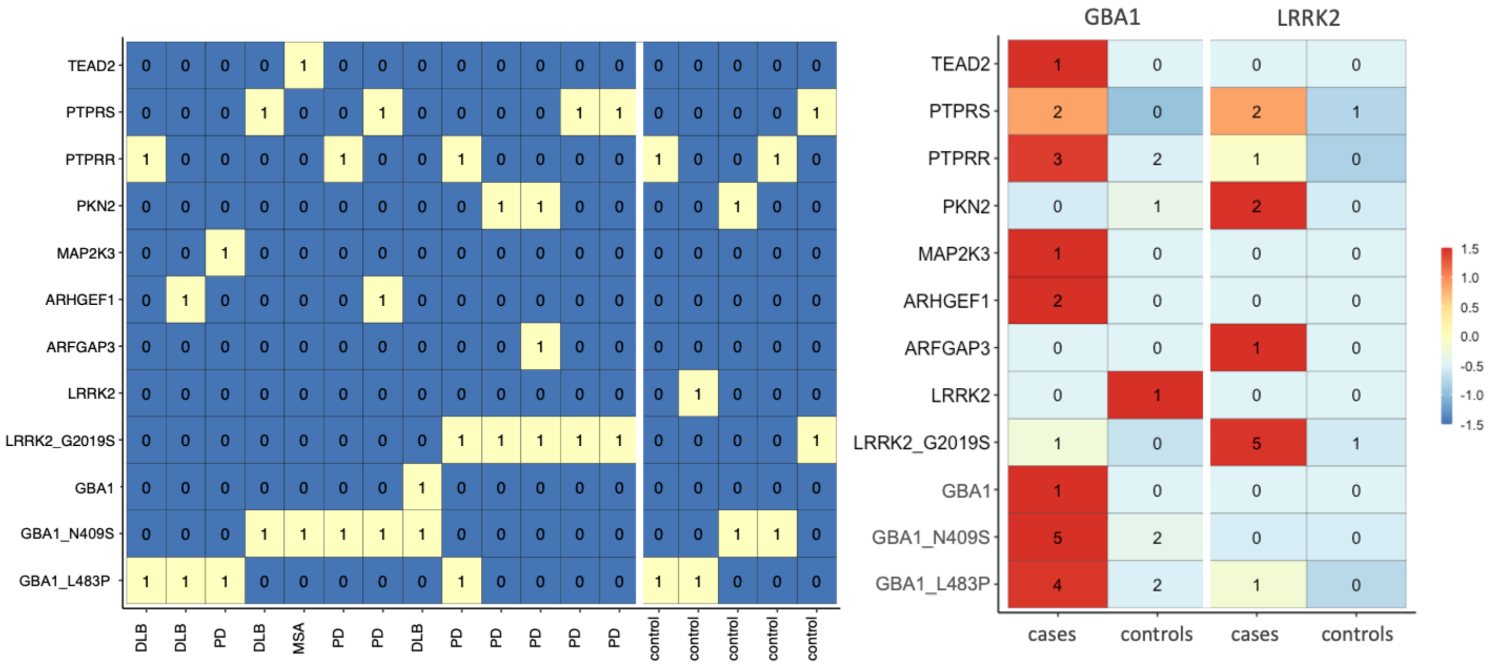
TEAD2
PTPRS
PTPRR
PKN2
MAP2K3
ARHGEF1
ARFGAP3
LRRK2
LRRK2_G2019S
GBA1
GBA1_N409S
GBA1_L483P
DLB
PD
MSA
control
GBA1
LRRK2
cases
controls
1.5
1.0
0.5
0.0
-0.5
-1.0
-1.5

**Supplementary Figure 2. Carriers of *GBA1* N409S (rs76763715), *GBA1* L483P (rs421016), and *LRRK2* G2019S (rs34637584).** Numbers represent total number of samples with a rare (MAF <= 1%) nonsynonymous variant in a stop gene. For case samples that harbor both a *GBA1* and a *LRRK2* mutation, ratio of carrier cases to all cases is shown scaled by genes *GBA1* and *LRRK2*. Similarly, control samples that harbor mutations in both the genes, ratio of carriers to all MGRB controls in shown scaled by the genes.

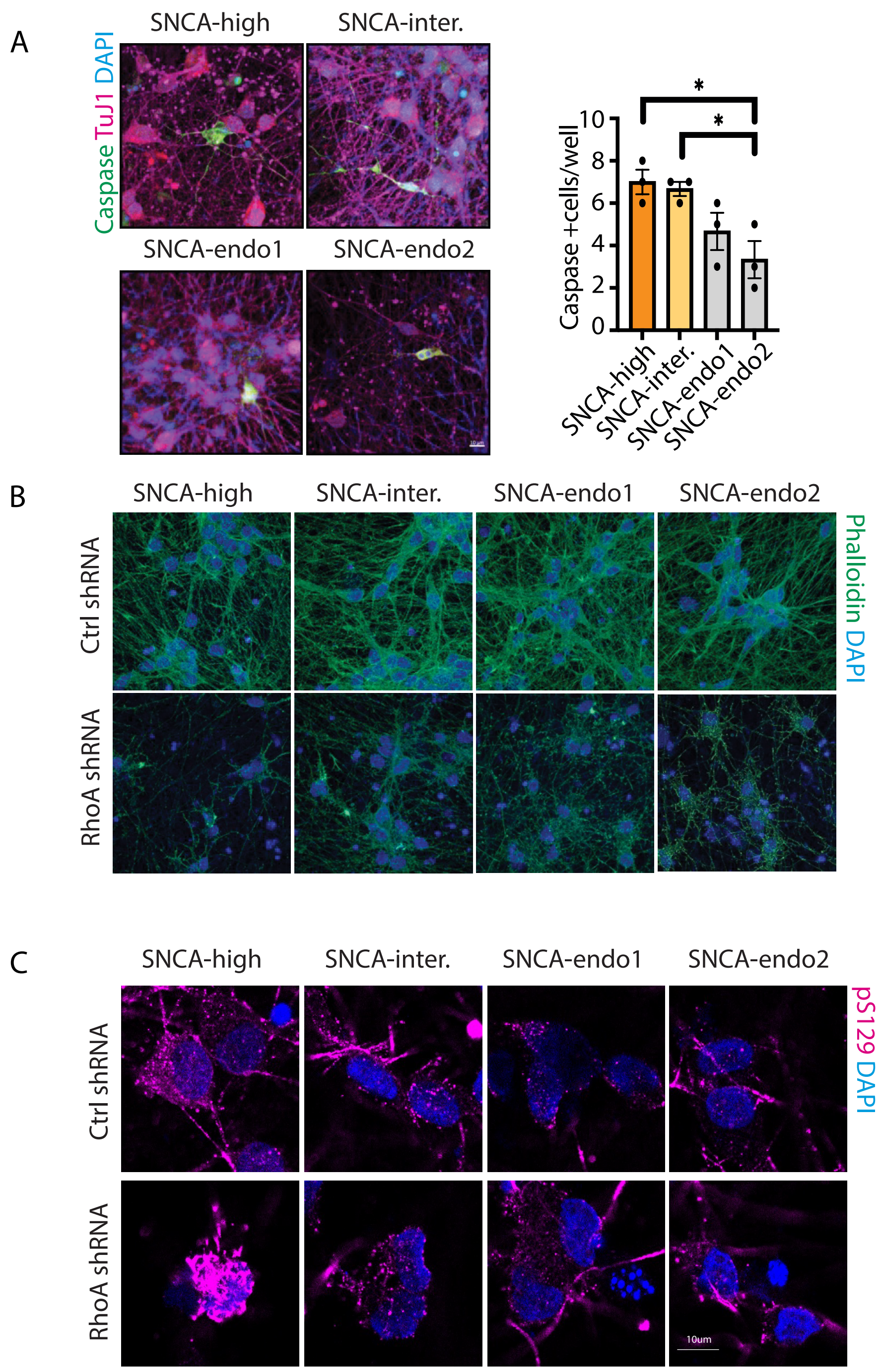
A
SNCA-high
SNCA-inter.
SNCA-endo1
SNCA-endo2
Caspase TuJ1 DAPI
Caspase +cells/well
10
8
6
4
2
0
*
*
SNCA-high
SNCA-inter.
SNCA-endo1
SNCA-endo2
B
SNCA-high
SNCA-inter.
SNCA-endo1
SNCA-endo2
Ctrl shRNA
RhoA shRNA
Phalloidin DAPI
C
SNCA-high
SNCA-inter.
SNCA-endo1
SNCA-endo2
Ctrl shRNA
RhoA shRNA
pS129 DAPI
10um

**Supplementary Figure 3: Significantly higher Caspase+ cells in *SNCA* overexpression neurons. RhoA downregulation phenocopies *SNCA* overexpression.**
(A) Sample confocal images of DIV28 CiS neurons immunostained with Caspase, TuJ1 and DAPI, showing significantly increased Caspase+ cells in SNCA-overexpression CiS neurons. Values represent mean ± s.e.m. (*$p < 0.05$; one-way ANOVA).
(B-C) Sample confocal images of CiS neurons transduced with control or RhoA shRNA. DIV14 neurons immunostained with (B) Phalloidin, DAPI; (C) pS129, DAPI.

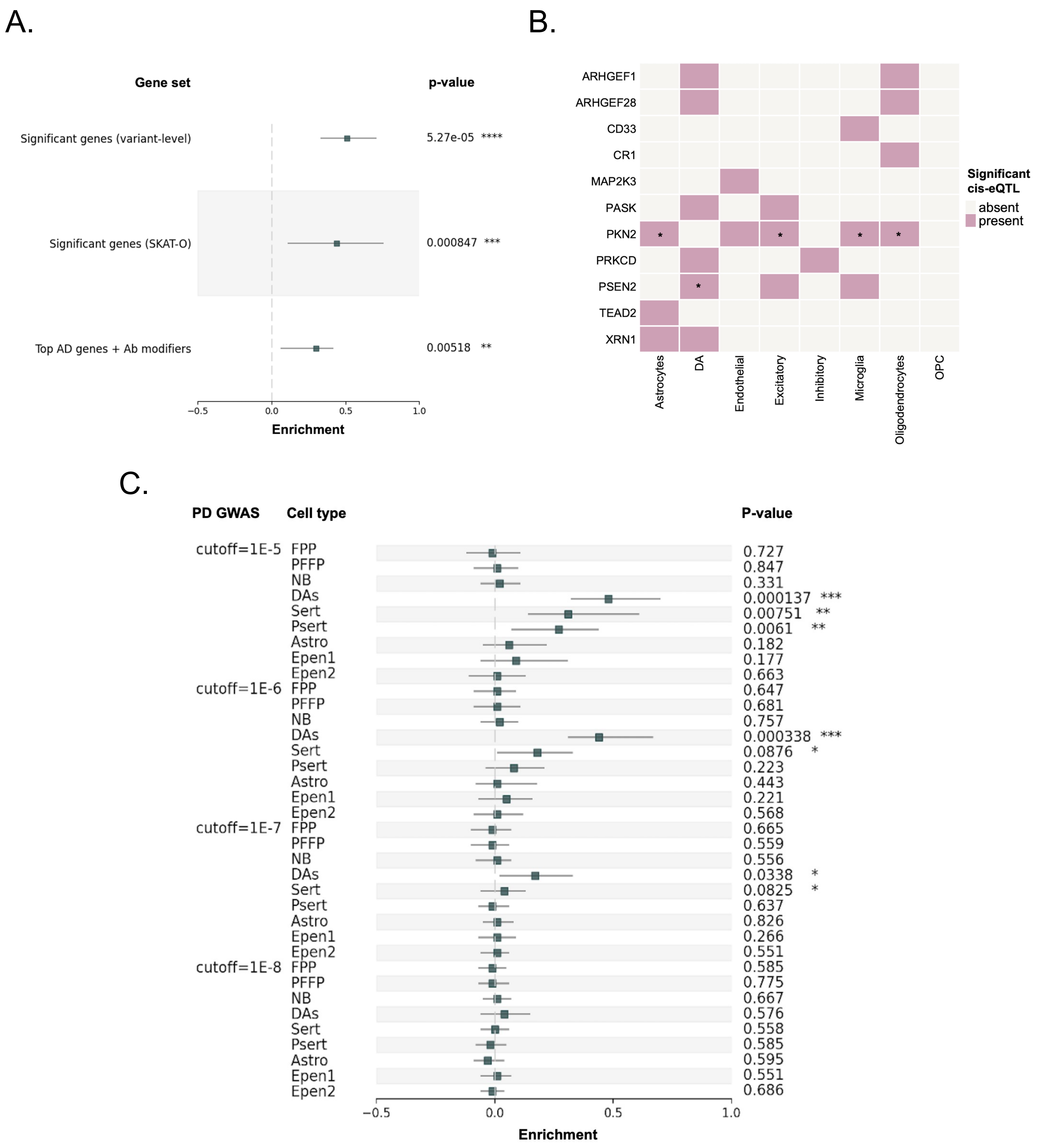
A.
Gene set
p-value
Significant genes (variant-level)
5.27e-05 ****
Significant genes (SKAT-O)
0.000847 ***
Top AD genes + Ab modifiers
0.00518 **
−0.5
0.0
0.5
1.0
Enrichment
B.
ARHGEF1
ARHGEF28
CD33
CR1
MAP2K3
PASK
PKN2
PRKCD
PSEN2
TEAD2
XRN1
Astrocytes
DA
Endothelial
Excitatory
Inhibitory
Microglia
Oligodendrocytes
OPC
Significant cis-eQTL
absent
present
C.
PD GWAS
Cell type
P-value
cutoff=1E-5
FPP 0.727
PFFP 0.847
NB 0.331
DAs 0.000137 ***
Sert 0.00751 **
Psert 0.0061 **
Astro 0.182
Epen1 0.177
Epen2 0.663
cutoff=1E-6
FPP 0.647
PFFP 0.681
NB 0.757
DAs 0.000338 ***
Sert 0.0876 *
Psert 0.223
Astro 0.443
Epen1 0.221
Epen2 0.568
cutoff=1E-7
FPP 0.665
PFFP 0.559
NB 0.556
DAs 0.0338 *
Sert 0.0825 *
Psert 0.637
Astro 0.826
Epen1 0.266
Epen2 0.551
cutoff=1E-8
FPP 0.585
PFFP 0.775
NB 0.667
DAs 0.576
Sert 0.558
Psert 0.585
Astro 0.595
Epen1 0.551
Epen2 0.686
−0.5
0.0
0.5
1.0
Enrichment

**Supplementary Figure 4. Enrichment of expression quantitative trait loci (eQTL) near SKAT-O significant AD genes and Aβ toxicity modifiers.**

(A) Enrichment of variant-level and gene-level significant genes from rare variant analysis as well as the set of top AD genes and Aβ modifiers among genes with e/sQTLs in both GTEx Brain and psychENCODE datasets. Confidence intervals were empirically estimated through random sampling of N (N=number of member genes in target set), permuting 1,000 times.

(B) Presence of significant cell-type specific cis-eQTLs in different brain cell types near the top AD genes and Aβ modifiers. We observed enrichment of cell-type-specific cis-eQTLs in DA neurons near PSEN2 and PKN2 was enriched for cis-eQTLs in astrocytes, oligodendrocytes, microglia, and excitatory glutamatergic neurons (hypergeometric p-value < 0.05).

(C) Enrichment of cell-type specific cis-eQTLs near top AD genes and Aβ modifiers in different iPSC-derived cell types towards a mid-brain neuronal fate and their link to PD GWAS. Enrichment analysis was performed using different GWAS p-value thresholds < $1x10^{-8}$. The AD genes and Aβ modifiers set was enriched in DA neuron-specific regulatory variants with suggestive links to PD at different cutoffs from $1x10^{-5}$ to $1x10^{-7}$. Confidence intervals were empirically estimated through random sampling of N (N=Number of genes in the target gene set), permuting 1,000 times. Here, FPP: Floor plate progenitors, PFPP: Proliferating floor plate progenitors, NB: Neuroblasts, DAs: Dopaminergic neurons, Serts: Serotonergic-like neurons, PSerts: Proliferating Serotonergic-like neurons, Astro: Astrocyte-like, Epen1: Ependymal-like 1, Epen2: Ependymal-like 2.

A.

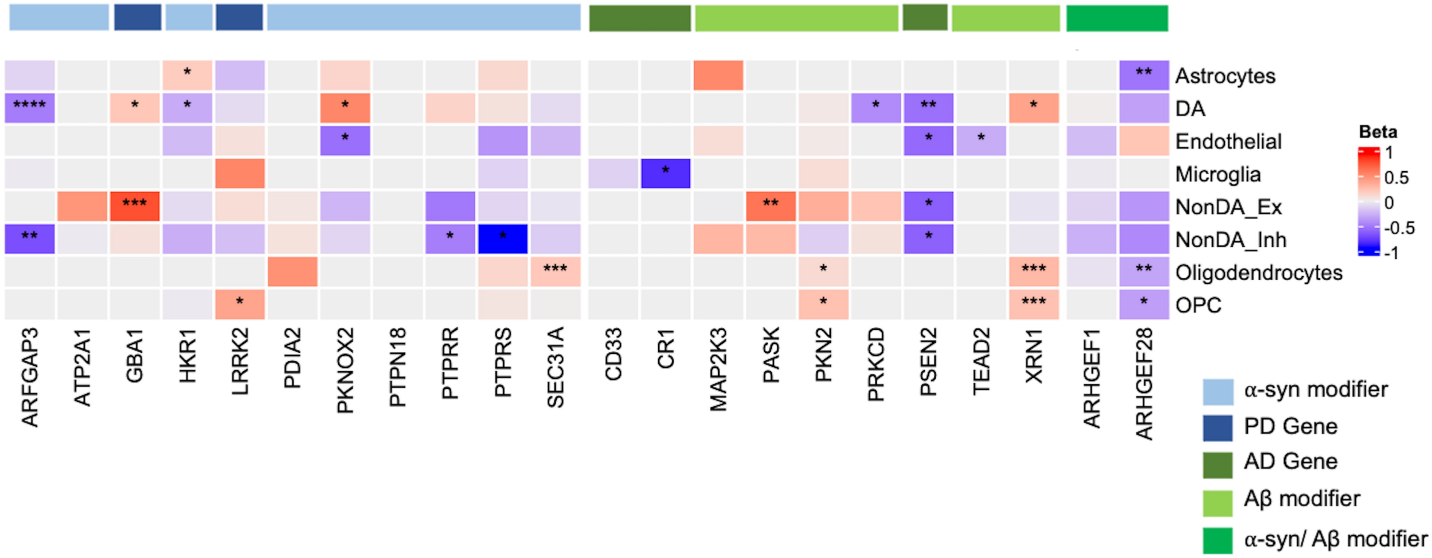
Astrocytes
DA
Endothelial
Microglia
NonDA_Ex
NonDA_Inh
Oligodendrocytes
OPC
ARFGAP3
ATP2A1
GBA1
HKR1
LRRK2
PDIA2
PKNOX2
PTPN18
PTPRR
PTPRS
SEC31A
CD33
CR1
MAP2K3
PASK
PKN2
PRKCD
PSEN2
TEAD2
XRN1
ARHGEF1
ARHGEF28
Beta
1
0.5
0
-0.5
-1
α-syn modifier
PD Gene
AD Gene
Aβ modifier
α-syn/ Aβ modifier

B.

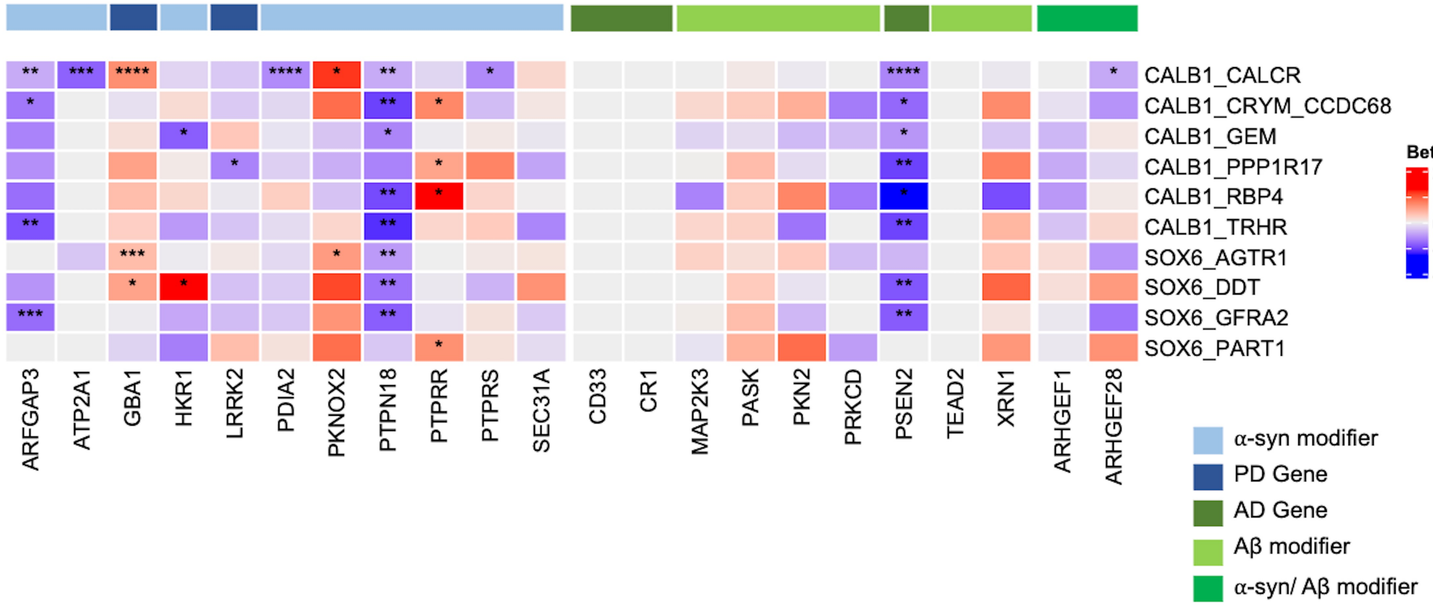
CALB1_CALCR
CALB1_CRYM_CCDC68
CALB1_GEM
CALB1_PPP1R17
CALB1_RBP4
CALB1_TRHR
SOX6_AGTR1
SOX6_DDT
SOX6_GFRA2
SOX6_PART1
ARFGAP3
ATP2A1
GBA1
HKR1
LRRK2
PDIA2
PKNOX2
PTPN18
PTPRR
PTPRS
SEC31A
CD33
CR1
MAP2K3
PASK
PKN2
PRKCD
PSEN2
TEAD2
XRN1
ARHGEF1
ARHGEF28
Beta
2
1
0
-1
-2
α-syn modifier
PD Gene
AD Gene
Aβ modifier
α-syn/ Aβ modifier

**Supplementary Figure 5. Differential expression of our significant genes across different CNS cell types and different subtypes of DA neurons in PD/DLB brains.**
(A) Heatmap showing PD-associated transcriptional changes of the significant 22 genes from our rare variant analysis in post-mortem PD/DLB brains vs controls in different CNS cell types. Cell-type specific differential expression analysis was performed using MAST's discrete hurdle model. Transcriptome-wide significance is reported using a BH-corrected p-value cutoff of 0.1 (denoted by *s; *: adj-p < 0.1, **: adj-p < 0.01, ***: adj-p < 0.001, ****: adj-p < 0.0001).
(B) Heatmap showing PD-associated transcriptional changes of top 22 in post-mortem PD/DLB brains vs controls in different subtypes of DA neurons in the midbrain. Cell subtype specific differential expression analysis was performed using MAST's discrete hurdle model. Transcriptome-wide significance is reported using a BH-corrected p-value cutoff of 0.1 (denoted by *s; *: adj-p < 0.1, **: adj-p < 0.01, ***: adj-p < 0.001, ****: adj-p < 0.0001).

**A.**

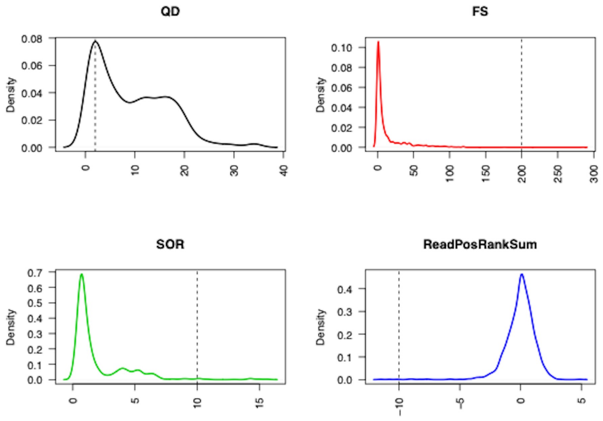


*Hard filters (indels)*

**B.**

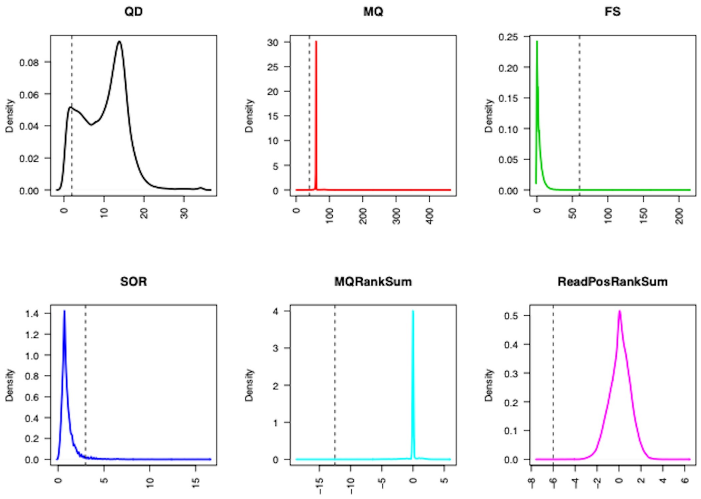


*Hard filters (SNPs)*

**Supplementary Figure 6. Quality metrics used for variant-level filtering in our targeted exome sequencing dataset. (**A) GATK hard filters used for indels. (B) GATK hard filters for SNVs.

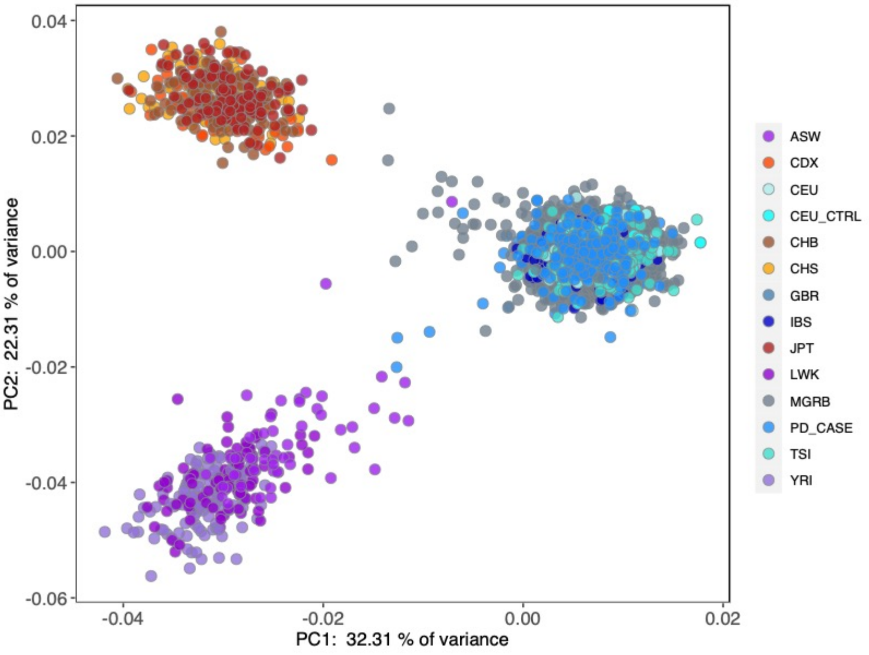

PC1: 32.31 % of variance
PC2: 22.31 % of variance
0.04
0.02
0.00
-0.02
-0.04
-0.06
-0.04
-0.02
0.00
0.02
ASW
CDX
CEU
CEU_CTRL
CHB
CHS
GBR
IBS
JPT
LWK
MGRB
PD_CASE
TSI
YRI


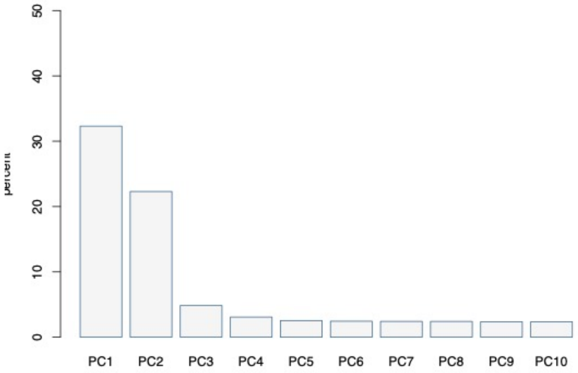

0
10
20
30
40
50
PC1
PC2
PC3
PC4
PC5
PC6
PC7
PC8
PC9
PC10

**Supplementary Figure 7. Principal component analysis of synucleinopathy cases in our targeted exome sequencing cohort, HapMap CEU controls and MGRB controls with labeled 1000 Genomes samples.** Variance explained by each PC is shown on the right.

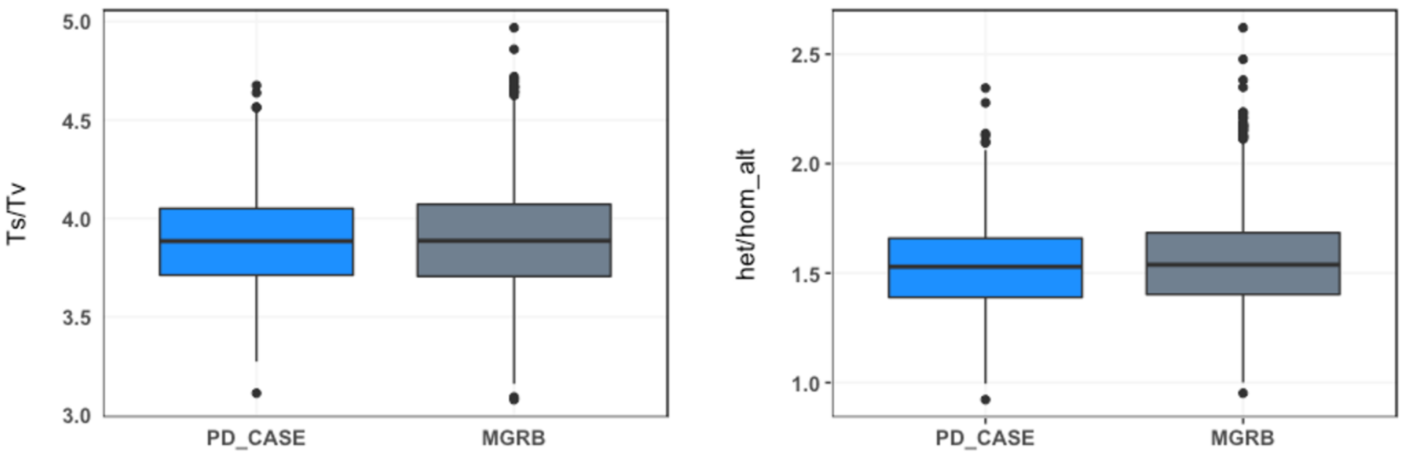

Ts/Tv
5.0
4.5
4.0
3.5
3.0
PD_CASE
MGRB
het/hom_alt
2.5
2.0
1.5
1.0
PD_CASE
MGRB

**Supplementary Figure 8. Distribution of Ts/Tv and Het/Hom ratio in our targeted exome sequencing dataset.**
(A) Boxplot of Ts/Tv ratios in cases vs MGRB controls showed no significant difference (Wilcoxon p=0.5854).
(B) Boxplot of heterozygosity per sample, measured as the ratio between heterozygous SNVs and homozygous alternate SNVs in cases and MGRB showed no significant differences between the two groups (Wilcoxon p=0.13).

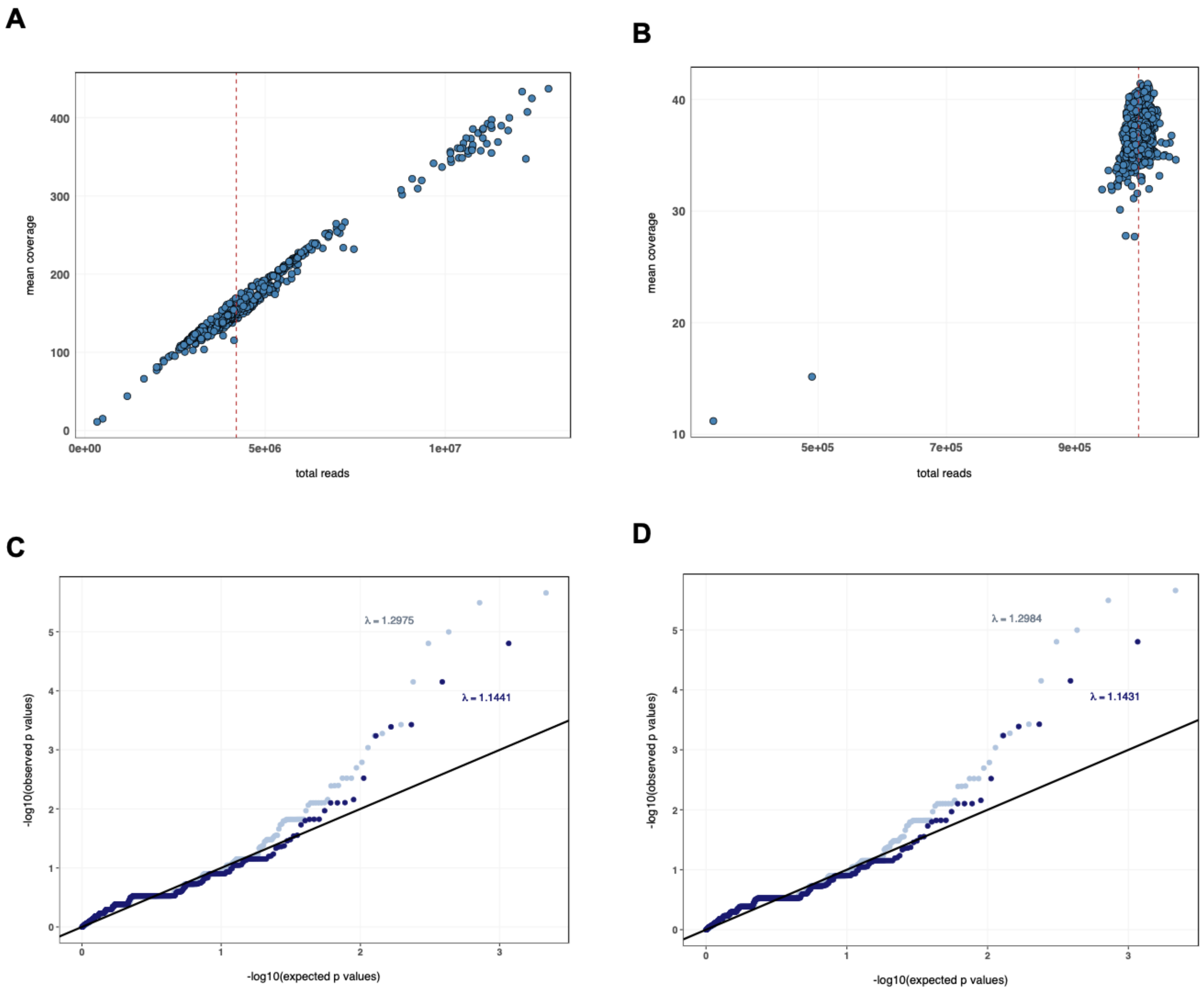
A
mean coverage
total reads
0
100
200
300
400
0e+00
5e+06
1e+07
B
mean coverage
total reads
10
20
30
40
5e+05
7e+05
9e+05
C
λ = 1.2975
λ = 1.1441
-log10(observed p values)
-log10(expected p values)
D
λ = 1.2984
λ = 1.1431
-log10(observed p values)
-log10(expected p values)

**Supplementary Figure 9. Downsampling experiment to ensure the cause of genomic inflation.**

(A) Mean coverage per sample in the target regions versus total uniquely mapped reads.
(B) Mean coverage after downsampling each sample to 1 million reads. The sample with ~11X coverage did not pass variant and sample level quality filters and was excluded from the analysis.
(C) Quantile-Quantile plots from a Fisher's Exact test of nonsynonymous variants detected using all reads.
(D) Quantile-Quantile plots from a Fisher's Exact test of nonsynonymous variants detected after downsampling.
